# Ancient amino acid sets enable stable protein folds

**DOI:** 10.1101/2025.10.29.685319

**Authors:** V. G. Giacobelli, S. Andersson, P. Srb, T. Neuwirthová, Z. Ruszová, J. Marhoul, S. Pšenička, A. Knetl, L. Bednárová, V. Veverka, K. Hlouchová, I. André

## Abstract

Early proteins likely arose from a chemically limited set of amino acids available through prebiotic chemistry, raising a central question in molecular evolution: could such primitive compositions yield stable, functional folds? Using *de novo* design, we constructed three ancient protein architectures using a reduced, evolution-inspired alphabet of ten amino acids, e.g. lacking all basic and aromatic residues. The resulting structures adopted their intended topologies and showed exceptional resistance to thermal and chemical denaturation. Computational simulations further revealed that proteins built from this restricted alphabet were as mutation-resilient as those using all twenty canonical residues. Besides their evolutionary implications, our results provide a foundation for minimalist protein design and generation of simplified, robust systems in minimal cell engineering.

## Main Text

The canonical set of 20 amino acids forms the foundation of modern proteins, reflecting a long evolutionary history of selection and optimization. This biological alphabet has remained largely conserved since the Last Universal Common Ancestor (LUCA), exhibiting only minor variations across species and time^1,2^. It has been proposed that the canonical set is near-optimal, balancing chemical diversity with the constraints of the genetic code^3^. However, the question of why life settled on precisely these 20 amino acids remains one of the most intriguing puzzles of molecular evolution^4–6^.

Pre-LUCA life likely relied on a reduced subset of amino acids. Notably, prebiotic chemistry could only generate a limited set of canonical “early” amino acids—primarily small, polar, and aliphatic residues such as the canonical Gly, Ala, Asp, Glu, Val, Ser, Ile, Leu, Pro, and Thr—lacking both aromatic and basic side chains^7^. It has long been questioned whether proteins built exclusively from this early canonical set could fold into stable, functional structures, given the restricted physicochemical diversity^8^. Computational and experimental studies suggest the early set possesses inherent propensities for secondary and tertiary structure formation^9–16^ and sequences composed of this acidic alphabet are also substantially more soluble compared with the “full” alphabet^10,11,17,18^. Despite these leads, no purely early-alphabet folds have yet been designed and characterized.

Traditionally, protein design initiatives are based on the biology-inspired amino acid alphabet. Rare efforts in engineering proteins with reduced amino acid alphabets have explored minimal alphabets that typically aim to preserve key chemical properties while simplifying design complexity^8,19–23^. However, most reduced-alphabet proteins reported to date exhibit compromised stability or solubility, limiting their practical applications.

Here, we leverage modern protein design tools, RFdiffusion and ProteinMPNN, to design proteins restricted to the prebiotic early amino acid set. By populating three ancient protein folds with only early amino acids, we successfully test the hypothesis that the early alphabet can support stable, compact protein structures. Biophysical characterization of these designed proteins demonstrates robust folding, high solubility, and ease of expression. Besides its potential implications for pre-LUCA life, evolution-guided restriction of the protein alphabet presents a possible protein design strategy towards increased protein solubility and significantly decreased metabolic costs.

## Results

### Design of Ancient Protein Folds Using a Restricted Amino Acid Alphabet

This study aims to assess whether proteins with ancient folds can adopt stable, well-defined structures when constructed from a restricted amino acid alphabet proposed to have been available on prebiotic Earth. The chosen ancient folds were taken from a phylogenetic study carried out by Caetano-Anollés *et al*., which includes a phylogenomic tree describing the evolution of 776 folds defined by the SCOPe database^24^. The 9 most ancient folds, which appeared at the base of the tree and are widespread in metabolism across the kingdom of life, were chosen. These folds were, in order from oldest to newest: P-loop containing nucleoside triphosphate hydrolases, DNA/RNA-binding 3-helical bundle, TIM α/β-barrel, NAD(P)-binding Rossman-fold domains, Ferredoxin-like, Flavodoxin-like, Ribonuclease H-like motif, OB-fold, and S-adenosyl-L-methionine-dependent methyltransferases. The phylogenetic relationships in the tree of these nine ancient folds were congruent with the relationships found in other global trees^24^. Sequences and structures of the 9 ancient folds were extracted from the SCOPe database^25^ based on their fold classification, removing redundant sequences with more than 40% identity. We focused on sequences between 50 and 100 residues, i.e. protein size that would be convenient for structural characterization by NMR. The overall distribution of domain lengths for each fold before and after filtering can be seen in Fig. S1. After filtering, there were 244 entries for DNA/RNA-binding 3-helical, 230 entries for Ferredoxin, 81 entries for OB-fold, 8 entries for Ribonuclease H-like motifs, and 6 entries for P-fold hydrolase.

While it would be possible to redesign the existing structures with new sequences, the goal of this study was to design *de novo* proteins with the same fold as the original structure, but with a new structure and sequence. This would enable synergy between backbone conformation and protein sequence. To achieve this goal, the capabilities of the RFdiffusion method^26^ to condition *de novo* design on a fold were utilized. For each selected entry in the fold database, a scaffold was generated and used as a constraint to generate *de novo* structures with the same fold as the template, with 10 designs being created for each template protein. Sequence design was carried out using ProteinMPNN, which has been shown to be highly successful in designing well-folded sequences in combination with RFdiffusion^27^. The allowed amino acids were taken from Higgs *et al*., who correlated calculated ΔG values of formation for the amino acids under prebiotic conditions with the observed amino acid abundances^7^. The 10 amino acids expected to have emerged first were selected for our restricted alphabet and included G, A, D, E, V, S, I, L, P, and T. To obtain sequences with only these amino acids, a penalty was applied to the other 10 amino acids during sequence design with ProteinMPNN. This approach successfully restricted the amino acid composition to only the desired 10 amino acids. For each system, 20 sequences were generated. The best sequence for each structure was chosen based on the score provided by ProteinMPNN.

### Computational Validation of *De Novo* Designed Ancient Folds

A first validation of the generated structures was carried out with the models generated with RFdiffusion by comparing the desired and actual secondary structure content. At this stage, only the presence of the expected number of alpha helices and beta-strands was considered. During design, 5 structures were of too poor quality to create a secondary structure scaffold, which meant these were excluded from further generation. Out of a total of 5640 generated structures, 1082 had the correct secondary structure. On a per-class basis, DNA/RNA-binding 3-helical had a 23% success rate, Ferredoxin-like had an 18% success rate, OB-fold had a 13% success rate, Ribonuclease H-like had a 9% success rate, and lastly, P-fold Hydrolase had a 10% success rate.

The structural quality of the sequences was initially screened using ESMFold, due to its high computational efficiency and speed ^28^. pLDDT values from ESMFold were used as a quality metric, but the most important criteria were secondary structure and topology matching the template structure. Out of the 1082 designs validated in this way, 596 had the desired secondary structure and topology. At this stage, there were no valid P-fold Hydrolase structures, and the fold was thus excluded from further testing (Fig. S2). Three candidates were selected from each remaining fold class based on pLDDT: DNA/RNA-binding 3-helical, Ferredoxin, OB-fold, and Ribonuclease H-like motif.

A final validation with AlphaFold2^29^ was used to select sequences for experimental validation. All chosen designs were predicted as their respective fold by FoldSeek^30^, except for Ribonuclease. All the top matches for the generated Ribonuclease structures were unindexed by SCOPe. However, the majority of top matches had the same class, architecture, and topology for their CATH classification as the original structures and were thus considered a match in combination with manual verification of the fold^31^.

Application of ProteinMPNN on RFdiffusion templates can sometimes produce low complexity segments in sequences consisting of repeats of a single amino acid, in particular, alanine in alpha-helical regions. These features can result in issues in protein expression and NMR analysis. Low-complexity regions were manually edited by the introduction of alanine to glutamic acid substitutions. These variants had the same or improved pLDDT compared to the automatically designed sequences. Generally, a pLDDT of above 80 for AlphaFold predictions was considered a good structure, but these requirements had to be relaxed for both the OB-fold and Ribonuclease H-like fold, which had pLDDTs in the 60s and 70s. However, it is important to keep in mind that ESMFold or AlphaFold have not been trained on sequences with a restricted amino acid alphabet. This means that while a high pLDDT is promising, it does not mean that a sequence with a low pLDDT cannot fold^32^. The entire design workflow, along with final candidates chosen for experimental characterization is shown in Fig. 1A.

**Fig. 1.**
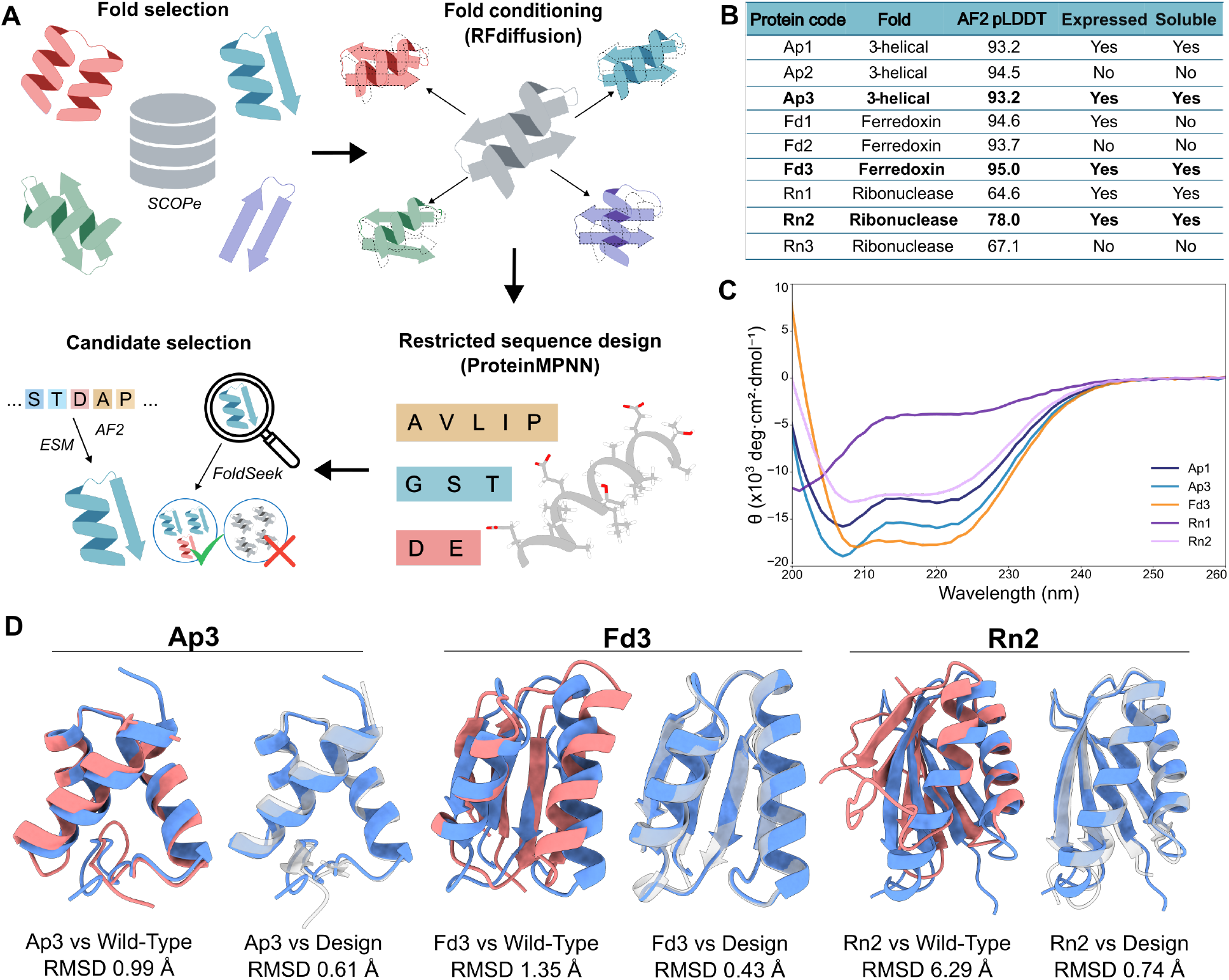
Experimental characterization and structural analysis of designed ancestral proteins. (A) Schematic representation of the design workflow for ancestral proteins. (B) Summary table reporting general features of the selected designs and their expression outcomes; proteins successfully expressed in the laboratory are indicated in bold. (C) CD spectra of the purified ancestral proteins. (D) Structural comparisons of Ap3, Fd3, and Rn2, respectively. Experimentally determined NMR structures are shown in blue, the corresponding wild-type templates in red, and the designed models from RFdiffusion in grey.

### Experimental Validation of *De Novo* Designed Ancient Folds

Out of nine designed constructs, five (Ap1–3, Fd3, and Rn1–2) were successfully expressed in *E. coli* BL21(DE3) as soluble proteins (Fig. 1B). These were purified and subjected to initial screening of structural properties using circular dichroism (CD) (Fig. 1C) and one-dimensional nuclear magnetic resonance (1D NMR) spectroscopy (Fig. S3).

CD spectroscopy revealed distinct secondary structure profiles for 4/5 expressed proteins. Ap1 and Ap2 displayed characteristic minima at 208 and 222 nm, consistent with a predominant α-helical content, while Fd3 and Rn2 showed spectral features typical of α/β proteins. In contrast, Rn1 exhibited a minimum at ∼200 nm, indicative of a predominantly disordered conformation lacking defined secondary structure. Among the four proteins with confirmed secondary structure content, Ap3, Fd3, and Rn2 displayed well-dispersed amide proton resonances (region of 7.5–9.5 ppm) in 1D NMR, indicative of a well-folded core. These three proteins were thus selected for further characterization and NMR studies aimed at determining their atomic-resolution structures.

### Three distinct folds constructed from ancient amino acids recapitulate the template architecture

Solution-state NMR spectroscopy revealed that all three ancient-sequence variants folded into well-defined and compact tertiary structures. For each construct, extensive chemical shift assignments, NOESY-derived distance restraints, and dihedral angle constraints enabled the calculation of converged structural ensembles with low backbone RMSDs relative to their respective wild-type templates (Fig. 1D).

The solution structure of protein Ap3 revealed a rigid Z-binding domain fold, characterized by a compact α/β architecture comprising three α-helices arranged around a central twisted β-sheet, forming a wedge-like structure. The well-defined hydrophobic core is maintained by a network of eight leucines, two isoleucines, four valines, and four alanines. Fourteen solvent-exposed glutamic acid residues and a single aspartic acid define an overall acidic character to the molecular surface. Overall, the structural ensemble displays minimal conformational variability, restricted to three residues at both the N- and C-termini.

The Rn2 protein adopts an RNase H–like fold with a mixed α/β architecture typical of nucleic acid–processing enzymes. Unlike the canonical structure, it contains a central four-stranded rather than five-stranded β-sheet, flanked on both sides by α-helices. Hydrophobic core stabilization is primarily mediated by six of the eight leucine residues located along the α-helices, together with three isoleucines and eight alanines distributed throughout the fold. Additional stabilizing contacts are provided by valine side chains, with seven of the nine residues located along the β-sheet strands. Negatively charged glutamic and aspartic acid residues are enriched within the solvent-exposed α-helical regions. The structural ensemble converged with conformational variability only observed in the N- and C-terminal segments. A minor second set of NMR signals was detected for backbone resonances of residues within α-helices and connecting loops. However, the NOE patterns in the 3D ^15^N-edited NOESY spectrum closely matched those of the major form.

The protein Fd3 adopts a compact, well-ordered fold comprising a central β-sheet flanked by two α-helices, forming a canonical nucleic acid–binding architecture commonly observed in viral proteins. An extensive network of hydrophobic residues stabilizes the core, including seven leucines, six isoleucines, six valines, and four alanines. Negatively charged glutamic and aspartic acid residues are predominantly clustered around two solvent-exposed helical regions. As in the other designs, conformational variability is restricted to the N- and C-terminal regions. The NMR spectra also reveal a second set of backbone resonances for residues surrounding proline 46, consistent with the presence of a cis peptide bond. This heterogeneity is restricted to backbone resonances and is not accompanied by a corresponding second set of side-chain resonances for the affected residues, suggesting that this region does not adopt a substantially distinct alternative conformation.

Overall, the quality of the NMR spectra and the precision of the structural ensembles were comparable to those typically obtained for modern proteins of similar size and topology, highlighting the intrinsic foldability and structural order of sequences built from a highly reduced chemical alphabet. Similarly, the experimental structures are highly similar (with RMSD of 0.4-0.7 Å) to the designed structure, reflecting high performance of the design pipeline with the reduced alphabet.

### One of the designed folds has copper-ion binding affinity

The Fd3 fold was the only structure among those examined that was derived from a metal-binding domain. Although metal-binding functionality was not considered during the design process, we assessed the affinity of Fd3 for several divalent cations (Mg^2+^, Zn^2+^, Fe^2+^, and Cu^2+^). Of these, only Cu^2+^ bound detectably, with an estimated dissociation constant (Kd) of ∼100 µM. The binding site comprises three glutamic acid residues (E16, E19, and E20), which coordinate the Cu^2+^ ion on the solvent-exposed surface of an α-helix (Fig. S4D). The addition of Cu^2+^ ions resulted in a substantial attenuation of resonance intensities in two-dimensional NMR spectra, consistent with direct metal ion – protein interactions (Fig. S4A-C). These data establish that polypeptides composed of an ancient subset of amino acids possess inherent metal-binding capacity. This property expands the chemical potential of minimal protein scaffolds, enabling them to generate coordination environments akin to those found in modern metalloproteins. By leveraging transition metals as prosthetic groups, such primitive sequences may have acquired catalytic functions beyond the intrinsic reactivity of their side chains^33–35^.

### Folds designed from ancient amino acids exhibit exceptional stability across diverse conditions

To evaluate the structural stability of the designed proteins, CD spectroscopy was performed under diverse conditions, covering a temperature range of 10–90 °C, a pH range of 4–8, and increasing concentrations of guanidine hydrochloride (GdnHCl) (Fig. S5A-I). Across the tested pH range, no significant alterations in the CD spectra were observed for most of the proteins, except for Rn2, which shows signs of aggregation or increased β-sheet formation at low pH (Fig. S5D-F). In addition, all three ancient proteins demonstrated relatively high thermal stability (Fig. 2A) (Fig. S5A-C). Only a modest reduction in ellipticity associated with α-helical content was detected, suggesting a partial loss of helical structure without major conformational rearrangement. Similarly, all three proteins also exhibited strong resistance to unfolding by GdnHCl (Fig. 2A) (Fig. S5G-I).

**Fig. 2.**
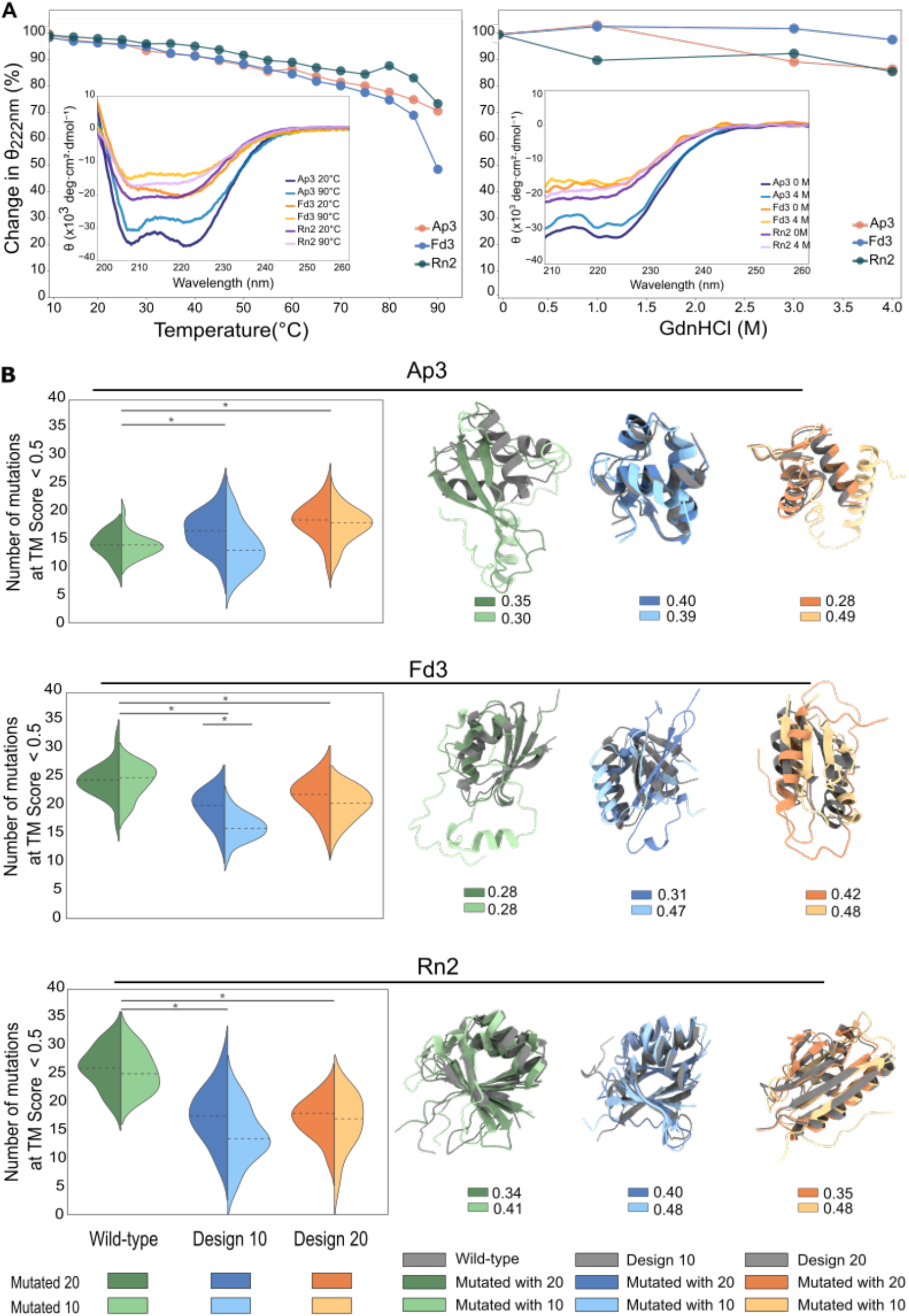
Stability and robustness of the designed folds. (A) Thermal and chemical stability of proteins Ap3, Fd3, and Rn2. Major plots show the change in mean residue ellipticity, [θ] (deg·cm^2^·dmol^−1^), at 222 nm as a function of temperature and guanidine HCl concentration (dots represent experimental data; the line is a guide to the eye). Minor plots compare CD spectra at 20 °C and 90 °C, as well as chemical denaturation in the absence or presence of 4 M guanidine HCl. (B) Mutational robustness of folds. Left: Violin plots show the number of mutations required for the TM-score to fall below 0.5 (indicating fold loss), based on 20 independent simulations per condition. Colors denote the amino acid alphabets used (see legend). Each half-violin represents one alphabet–protein combination, with consistent color coding throughout the figure. The dotted line marks the median. Except for Ap3, designs generated with the 20 amino acid alphabet are significantly less robust than their wild-type counterparts (P ≤ 0.01, Mann–Whitney U test, *). No significant differences are observed between the two alphabets. Simulations using the Ancient 10 alphabet show greater variability, with significant differences in all comparisons except for Rn2 design and Ap3 wild-type versus design. Two-alphabet simulations show no significant differences, except for the Fd3 Ancient 10 *de novo* design. Full statistical data and P-values are provided in Supplementary Data. Right: Gray structures depict either wild-type or *de novo* designs, overlaid with two mutated examples near the median mutation count for each alphabet (dark = All 20, light = Ancient 10). Numbers next to color boxes indicate TM-scores between each mutated structure and its corresponding starting model.

The pronounced resistance to thermal and chemical denaturation observed here contrasts with earlier studies of proteins incorporating ancient amino acid compositions^14,15,21^. In those cases, substitution toward ancient amino acids reduced hydrophobic core volume and introduced packing defects, even when mutations were designed to preserve structural integrity. Such instability has been attributed to the limited chemical diversity of the early amino acids, leading to the proposal that ancient proteins achieved stable conformations primarily under halophilic conditions. By contrast, the proteins examined in this work possess compact hydrophobic cores, similar to the wild-type template structures (Fig. S6). Two of the three designed folds have a higher fraction of hydrophobic residues compared to their template folds, which can partially explain their high stability (Fig. S7). In general, *de novo*-designed folds based on the ancient alphabet have a higher proportion of hydrophobic residues than folds designed with the complete set of amino acids.

To distinguish effects arising from the design procedure from those due to amino acid composition, we characterized a protein from a large library of sequence variants with the same composition as the computationally designed proteins, selected based on folding and solubility properties.

### Stability of ancient folds reflects design strategy rather than alphabet

We found that the designed folds were expressed at high levels in soluble form, without extensive optimization. Previous work has shown that high solubility is intrinsic to random sequences restricted to the ancient subset of the amino acid alphabet^10,17,18^. To compare such unevolved sequences with designed folds, we screened a library of 109-residue random proteins built from this alphabet for variants with high cytosolic expression in *E. coli* BL21(DE3) and compact predicted structures. Four sequences were selected based on (i) a dual-reporter assay for expression and (ii) low radius of gyration and solvent-accessible surface area values in ESMFold predictions. Two sequences were overexpressed at high yield, and one was purified for structural characterization. CD spectroscopy confirmed secondary structure elements, and ESMFold predictions indicated overall compactness, but 1D NMR spectra showed no evidence of a well-ordered core (Fig. S8A–B). Similar to molten globules–like proteins enriched in ancient residues^15^, this sequence gained secondary structural elements and thermal stability under high-salt conditions (Fig. S9A-E). This supports the hypothesis^16^ that proteins derived from ancient amino acids, enriched in acidic residues, would be stabilized in halophilic environments. While our design strategy yielded highly stable folds from these amino acids, the salt dependence observed here may have been critical for early proteins prior to evolutionary optimization.

### Reduced-alphabet designs retain robustness similar to full-alphabet designs

An important property of proteins is their ability to balance evolvability and structural stability. To assess whether this differs between reduced and full alphabets, the designs were repeated with the complete set of amino acids. In general, the designs of the complete set have higher predicted confidence scores, higher TM-scores/lower RMSD to the design model (Fig. S10). ESM3 and AlphaFold2 predictions of complete set designs showed 5.5-7.4% higher pLDDT scores, 9.2-13.9% higher TM-scores, and 31.4-35.0% lower RMSD compared to restricted set designs, while ESMFold predictions showed comparable performance between both sets.

The mutational robustness was then tested in silico for both the “ancient” and “full” designs along with wild-type templates. Only substitution mutations were considered, using either the “ancient” or “full” alphabet to assess their effects on fold robustness. Mutations were introduced randomly with equal probability for all amino acids. No selection was applied, and each protein–alphabet combination was simulated 20 times. Structures of each mutated sequence were predicted with ESMFold and scaffold tolerance was measured as the number of mutations at which the TM-score first dropped below 0.5, a threshold indicating a different fold^36^. A representative structure for the point where the TM-score fell below 0.5, indicating loss of the original fold, is shown in Fig. 2B. For Fd3 and Rn2, the wild-type variants are significantly more robust to mutations than any of the design folds. For Ap3, both “ancient” and “full” alphabet designs increased the fold robustness, but the Ap3 wild-type does not represent a fully natural sequence as it has already been previously mutated for increased stability^37^. The originally reported wild-type sequence exhibits essentially the same behaviour as our Ap3 wild-type construct, indicating that the observed robustness is intrinsic to the fold rather than a consequence of subsequent stabilizing mutations. (Fig. S11)

While the robustness of the “ancient” designs appears to be lower than their “full” counterparts (Fig. 2B), also seen in the median TM-scores in the trajectories, this difference is not statistically significant for 8 out of 9 compared pairs (Mann-Whitney U test, significant when p-value ≤ 0.01). In summary, the results suggest that folds with the “ancient” alphabet have similar structural robustness to the full set of 20 amino acids, implying comparable evolvability.

## Discussion

Early proteins likely emerged from a limited set of simple amino acids available through prebiotic chemistry, before the evolution of metabolism and templated synthesis expanded the molecular toolkit to the twenty canonical residues of modern life. Although many non-canonical amino acids could also form abiotically, their sizes and physicochemical properties closely resemble those of the prebiotically plausible canonical ones ^38^. This overlap implies that the first protein-like polymers were likely assembled from a limited chemical alphabet - simpler than that of extant proteins - raising a central question in molecular evolution: could such primitive compositions give rise to stable, folded structures capable of function? Our findings demonstrate that they could.

Using an early amino acid set thought to predate LUCA, we show that compact, well-defined folds can be encoded even within this chemically constrained, overall very acidic, alphabet. Of nine folds proposed to represent ancient protein topologies^24^, three consistently yielded successful designs, as assessed by target topology and ESMFold pLDDT predictions. The failure of the remaining six designs may reflect either constraints of the design pipeline used here or an intrinsic dependence of certain architectures on later-evolved residues. To explore the fold space available to a reduced alphabet represents an important challenge for future protein design efforts.

Intriguingly, metal binding emerged spontaneously in one of the characterized designs, suggesting that rudimentary functional properties might arise even from minimal alphabets. Exploring whether such scaffolds can also support more complex activities, such as ligand binding or catalysis, will be a key next step. It is assumed that the subsequent inclusion of “late” amino acids likely presented adaptive properties by expanding both the chemical functionality and the structural diversity of the protein universe^4,11,39^.

Proteins successfully designed with the 10 ancient amino acids are generally more hydrophobic than typical natural proteins, promoting very tight core packing. In the solved structures, their hydrophobic cores are fully occupied, and in two out of three cases they even exceed the packing density of their native counterparts. This indicates that the full space of side-chain geometries encoded with a canonical set is not required to densely pack the interior of proteins. This is supported by experiments with a *de novo* designed protein with the core consisting entirely of valine, retaining a very high thermal stability^40^. Consistent with these features, our ancient-alphabet proteins are extremely stable, resisting thermal unfolding. This agrees with prior work showing that the full set of 20 amino acids is not required to achieve a stable, well-packed protein core^22^. These findings suggest that the ancient scaffolds can be as robust from a stability standpoint as modern proteins, raising questions about how such robust folding will be balanced with the need for functional innovation in evolution.

Simulations and experiments show that extra stability enhances evolvability by allowing a protein to tolerate a wider range of functionally beneficial but destabilising mutations, and stabilising mutations can counterbalance the destabilising effects of adaptive mutations in nature^41,42^. Our mutational simulations suggest that *de novo* proteins designed with the ancient set were as robust to random mutations as those designed with the full set of twenty canonical amino acids. This comparable robustness implies that ancient scaffolds could be just as evolvable from a stability perspective. However, stability alone does not guarantee functional versatility; the ancient amino acid set samples a smaller chemical space than the canonical set, which may constrain the emergence of specific catalytic or binding functions despite similar robustness.

Beyond its evolutionary implications, our work shows that proteins composed of a restricted set of early, small amino acids can fold stably and express efficiently in *E. coli*, highlighting principles that extend beyond origin-of-life contexts. While modern AI-structure prediction methods have been trained on proteins with the full canonical alphabet, we find that they are capable of accurately predicting structures of the specific reduced evolution-inspired alphabet used here with confident pLDDT values. Because smaller and less chemically complex amino acids are typically less costly to synthesize, constraining the amino acid repertoire could represent a practical strategy for designing robust and energetically economical proteins^32^. This concept of using a smaller chemical space for protein design aligns with emerging efforts in synthetic and minimal cell engineering, where resource efficiency and functional simplicity are of key importance ^43^.

## Materials and Methods

### Selection of protein folds

Candidate protein superfamilies were chosen based on the 9 oldest folds identified by Caetano-Anollés *et al:* P-loop containing nucleoside triphosphate hydrolases, DNA/RNA-binding 3-helical bundle, TIM α/β-barrel, NAD(P)-binding Rossman-fold domains, Ferredoxin-like, Flavodoxin-like, Ribonuclease H-like motif, OB-fold, and S-adenosyl-L-methionine-dependent methyltransferases^24^. The SCOPe 2.08 ASTRAL sequence database was used to obtain the relevant sequences through the use of Regex filters to select only the relevant SCOPe superfamilies. The sequences, folds, SCOPe IDs, and lengths were stored and then filtered to only contain sequences with lengths between 50 and 100 residues. Chosen sequences were then mapped to a PDB database of SCOPe folds. If a sequence was a part of a larger protein, only the relevant fragment was extracted. All obtained sequences consisted of only one chain.

### Backbone structure generation

All obtained structures had to be processed into temporary smaller files as the fold conditioning generation, at the time, had issues with PDBs that had more than one model present, as was the case for NMR structures. To mitigate this issue, all structures had their first model and chain extracted into a separate file, which served as the basis for the fold conditioning. The secondary structure scaffolds for the fold conditioning were generated using the *make_secstruc_adj*.*py* script distributed by RFdiffusion. The generated constraint files were used as input to the RFdiffusion inference script to generate 10 designs per template.

### Sequence design

Sequence design was carried out on all structures obtained from RFdiffusion by using ProteinMPNN. First, a bias dictionary was created against the unwanted amino acids: N, K, Q, R, C, H, F, M, Y, and W. The chosen bias value was −10, which was found by testing different bias values and seeing when the unwanted amino acids stopped appearing. The *make_bias_AA*.*py* script distributed with ProteinMPNN was used to create biases, which were used as input to sequence design with ProteinMPNN. 20 sequences were designed for each template, and the output sequences were validated to ensure that no forbidden amino acids had been used in the generated sequences. Sequences designed with the complete alphabet (20 amino acids), used in the mutational trajectory analysis, were designed with the same pipeline.

### Candidate selection

First, a rough filtering was performed on the structures obtained from RFdiffusion. This was done by obtaining the secondary structure string for the template structure by using DSSP and simplifying it as follows:

1. Re-mapping the assignments to only denote alpha helices, beta sheets, and coils
2. Shortening the string by removing consecutive elements of the same type

The number of beta sheets and alpha helices of the template was then compared to all of its derivative structures. Order was not considered at this stage. The structures that passed this validation step had their best sequence predicted by ESMFold, where the best sequence was defined as having the lowest negative log-likelihood score output by ProteinMPNN. Once the structure was predicted, it underwent the same secondary structure validation as previously outlined. At this stage, all designs with the correct number of secondary structure elements were separated by their fold and ranked by their pLDDT, as obtained by ESMFold. Each sequence was manually inspected for quality, avoiding low complexity/high repeat sequences whenever possible. If only a part of a promising sequence was of poor quality, it was manually edited to improve quality. The only manual edits that were performed were in alanine-rich regions where, after manual inspections, some alanine residues were mutated into glutamic acid. The most promising sequences were then predicted using AlphaFold2 and underwent a final secondary structure validation. The pLDDT threshold was set to 80, except in the cases of OB-fold and Ribonuclease H-like, as they had fewer structures to choose from. Finally, the predicted structures were manually compared to the template to ensure the correct fold, along with searching the structures with FoldSeek. Both the RFdiffusion designs and the AlphaFold predictions were input to FoldSeek. The SCOPe classifications of the top FoldSeek matches were checked to see if the structure matched the correct folds. If a sequence passed all tests, it was chosen as a representative for its fold.

### Protein Selection and Expression Optimization

The coding sequences for Ap1–3, Fd1–3, Rn1–3 and Random10E, each appended with a C-terminal hexahistidine (6×His) tag, were synthesized by Thermo Fisher Scientific following codon optimization using IDT software. The synthesized genes were amplified *via* PCR and cloned into the pET24a(+) expression vector using *HindIII* and *XbaI* restriction sites. Sequence integrity of all constructs was verified by Sanger sequencing.

Recombinant protein expression was performed in *Escherichia coli* BL21(DE3) cells. Optimization experiments were conducted in 10 mL cultures of LB medium supplemented with kanamycin (50 µg/mL), varying isopropyl β-D-1-thiogalactopyranoside (IPTG) concentrations (0.5 or 1 mM), induction temperatures (25°C, 30°C, or 37°C), and post-induction incubation times (4 or 16 hours). Following incubation, 1 mL aliquots were harvested by centrifugation at 5,000 × *g* for 5 minutes at 4°C and resuspended in 100 µL of B-PER™ Bacterial Protein Extraction Reagent (Thermo Fisher Scientific). Lysates were clarified by centrifugation at 20,000 × *g* for 10 minutes at 4°C. Soluble and total protein fractions were analyzed via SDS-PAGE on 18% gels followed by Western blotting. Detection was carried out using a monoclonal anti-6×His HRP-conjugated antibody (Sigma-Aldrich) and Immobilon® Forte Western HRP substrate (Merck). Chemiluminescent signals were visualized using the Amersham Imager 600 (Cytiva). Optimal expression conditions were determined for each protein and applied in subsequent large-scale preparations.

### Protein Purification

For preparative-scale expression, freshly transformed *E. coli* BL21(DE3) cells were cultured overnight in 10 mL LB medium supplemented with kanamycin (50 µg/mL) at 30°C and 220 rpm. These pre-cultures were used to inoculate 500 mL of fresh LB medium (also containing kanamycin at 50 µg/mL), which was incubated at 37°C with agitation at 220 rpm. At an optical density at 600 nm (OD_600_) of approximately 0.5, IPTG was added to a final concentration of 0.5 mM to induce expression. Cultures were further incubated under the following conditions: 25°C for Ap3, Fd3, Rn2, Rn3 and Random 10E; 37°C for Ap1; for either 4 or 16 hours. Cells were harvested by centrifugation at 10,000 × *g* for 20 minutes at 4°C. Cell pellets were resuspended in B-PER (Thermo Fisher Scientific) at a ratio of 4 mL per gram of wet biomass. The lysis buffer was supplemented with DNase I (5 U/mL, Jena Bioscience) and lysozyme (100 µg/mL, Sigma-Aldrich). Lysis was conducted at room temperature for 20 minutes with gentle agitation (60 rpm), followed by clarification through centrifugation at 20,000 × *g* for 25 minutes at 4°C. Clarified lysates were applied to 0.5 mL Talon® Metal Affinity Resin (Clontech Laboratories, Inc.) pre-equilibrated with buffer containing 300 mM NaCl and 50 mM NaH2PO4 (pH 8.0). Binding was performed at 4°C for 30 minutes with continuous mixing. The resin was washed with buffer containing 300 mM NaCl and 50 mM NaH2PO4 (pH 7.0), and bound proteins were eluted with 2.5 mL of elution buffer (300 mM NaCl, 50 mM NaH2PO4, 250 mM imidazole, pH 7.0). For Fd3 constructs containing a TEV protease cleavage site, on-column cleavage was performed with 72 µg of TEV protease at 4°C for 15 hours in 1.5 ml of buffer containing 300 mM NaCl and 50 mM NaH2PO4 (pH 7.0). Eluted proteins were further processed using the same purification protocol as uncleaved variants. Size-exclusion chromatography (SEC) was carried out on a Superdex 75 10/300 GL column (Cytiva) equilibrated in one of the following buffers: PBS buffer (137 mM NaCl, 2.7 mM KCl, 10 mM Na2HPO4, 1.8 mM KH2PO4, pH 7.4). SEC fractions containing the target protein were pooled, concentrated, and dialyzed prior to further purification by anion exchange chromatography using a HiTrap CaptoQ 1 mL column (Cytiva), pre-equilibrated with 20 mM NaH2PO4 (pH 5.9). Proteins were eluted with a linear gradient of 0–100% buffer B (10 mM NaH2PO4, 1 M NaCl, pH 5.9). Final protein fractions were pooled, concentrated, dialyzed in PBS, and stored at –80°C. For structural studies by nuclear magnetic resonance (NMR), Ap3, Fd3, Rn2 and Random10E were expressed following the same protocol described above, except that cultures were grown in M9 minimal medium supplemented with isotopically labeled ^15N-ammonium chloride and ^13C-glucose for Ap3, Fd3 and Rn2 only, (Cambridge Isotope Laboratories, Inc.) to enable uniform labeling. Protein concentrations were quantified by amino acid analysis using a Biochrom 30+ Series Amino Acid Analyzer (Biochrom, UK).

### Construction of Dual Reporter Vectors

To generate the dual reporter (DR) system, the pET24a(+) (Addgene) plasmid was modified by inserting a synthetic expression cassette (Integrated DNA Technologies, IDT) comprising a Golden Gate-compatible cloning site, a FLAG tag, a ribosome binding site (RBS), and the fluorescent protein gene *mScarlet-I*. The cassette was ligated into pET24a(+) via *HindIII* and *XbaI* restriction sites. The plasmid pSEVA631 (Addgene) was digested with *PacI* and *SpeI*, and a DNA cassette containing the *IbpAp* promoter and an in-frame unstable GFP variant was inserted using sticky-end ligation (Sticky-End Ligation Master Mix, New England Biolabs). To confer ampicillin resistance, the gentamicin resistance cassette of pSEVA631 was replaced with a β-lactamase gene. This was achieved by digestion with *PpuMI* and *EaII*, followed by ligation with an ampicillin resistance cassette.

### Library Design and Cloning

A peptide library was designed using CoLiDe software (Tretyachenko et al., 2021): the “Random10Elib” library (representing early amino acids). Amino acid frequencies were determined from UniProt-derived distribution datasets. Two 199-nucleotide ssDNA oligonucleotides were synthesized (IDT). The dsDNA library was generated *via* a Klenow extension reaction. Single-stranded oligos were combined at a final concentration of 10 µM in 1X Klenow buffer, denatured at 95 °C, and annealed by slow cooling to 71 °C for 5 min. The reaction was then cooled to 25 °C, followed by the addition of 1 µM dNTPs and 5 U Klenow Fragment (DNA Polymerase I; NEB) for strand synthesis. The resulting dsDNA was purified using the Monarch PCR & DNA Cleanup Kit (NEB). Approximately 150 fmol of each dsDNA library was cloned into the modified pET24a(+) vector (containing the *mScarlet-I* cassette) via Golden Gate Assembly (NEB), using a 2:1 insert-to-vector molar ratio. The reaction was cycled for 5 min at 16 °C and 37 °C overnight. Assembly products were purified with the Monarch PCR & DNA Cleanup Kit and electroporated into *E. coli* 10G supreme electrocompetent cells (Duos). Transformants were plated on LB-kanamycin (50 µg/mL) plates, incubated overnight at 37 °C, and ∼300K cell colonies were pooled for plasmid isolation. The cell pellets were resuspended in cold PBS supplemented with kanamycin, and plasmid DNA was purified using the Zyppy Plasmid Miniprep Kit (Zymo Research).

### Co-transformation and Library sorting

For co-expression of the dual reporter system, 50 ng each of pSEVA631-IbpAp-GFP-Amp and pET24a(+)-Random10ELib-*mScarlet-I* were electroporated into 25 µL *E. coli* BL21(DE3) electrocompetent cells (Duos). Transformants were recovered in 1 mL SOC medium (NEB) at 37 °C for 1 hour with shaking and plated on LB agar containing ampicillin (100 µg/mL) and kanamycin (50 µg/mL). Colonies were harvested by washing the plates with cold PBS containing kanamycin and ampicillin. Approximately 1 million colonies were collected. Glycerol stocks were prepared at OD_600_ = 1.0 (20% v/v glycerol). Upon expression, the glycerol stocks were diluted with fresh LB to OD_600_ = 0.2, grown shaking at 37°C until reaching OD_600_=0.6 and cooled down. Protein expression was induced with 0.5 mM IPTG, followed by incubation at 25 °C for 16 h. Cells were harvested (5 min, 4,000 × g, 4 °C), washed three times with ice-cold PBS, and diluted 1:30 before analysis on a BD FACSAria Fusion flow cytometer equipped with a 70 μm nozzle and a 1.0 ND (neutral density) filter. The sample chamber was maintained at 4 °C, and collection tubes were kept at 30 °C. GFP fluorescence was detected using the 488 nm blue laser with a 525/20 bandpass filter (FITC channel), and mScarlet-I was detected using the 561 nm yellow-green laser with a 586/15 bandpass filter (PE channel). The SSC threshold was set to 300 to capture *E. coli* cell populations. Initial gating for size and shape was performed using SSC-A × FSC-A and FSC-H × FSC-A projections. Cells with high expression of both GFP and mScarlet-I (G1 gate) were identified using FITC × PE projections. Sorted cells were collected into rich recovery medium (LGC, Biosearch Technologies, Hoddesdon, UK) in 1.5 mL centrifuge tubes, plated onto LB agar supplemented with kanamycin and ampicillin, and incubated at 30 °C for 16 h. Colonies were subsequently collected for plasmid DNA extraction.

### High-throughput Sequencing

Plasmid pools recovered from FACS-sorted populations were used as templates for PCR amplification prior to next-generation sequencing (NGS). PCR reactions were carried out with Q5 High-Fidelity DNA Polymerase (NEB, Ipswich, MA) using 50 ng of plasmid DNA per 50 μL reaction and run for 11 cycles. Primers were designed to anneal to the pET24a(+)-Random10Elib-mScarlet-I backbone. Following purification, amplicon size distributions of selected samples were analyzed on an Agilent 2100 Bioanalyzer (Agilent Technologies, Santa Clara, CA). Sequencing libraries were prepared with the NEBNext Ultra II DNA Library Prep Kit for Illumina (NEB, Ipswich, MA). Library size and concentration were again verified by Bioanalyzer analysis. Samples were pooled and sequenced on an Illumina NextSeq platform using NextSeq 1000/2000 P1 reagents (600 cycles) and Illumina MiSeq v3 (600 cycles) in paired-end mode. Sequencing reads were merged, trimmed, and filtered to remove low-quality reads using *fastp*^44^. The reference library was generated by clustering the unsorted sequence pool at 99% nucleotide identity with *cd-hit*^45^. Sorted reads were mapped to this reference using the Burrows–Wheeler Aligner^46^ (BWA-MEM). Conversion to SAM format, sorting, and indexing were performed with SAMtools^47^. Read counts per variant were obtained with HTSeq^48^. Variants were selected for further analyses based on high post-sorting read counts and manual inspection of predicted structures generated by ESMFold.

### Circular Dichroism Spectroscopy

ECD spectra were recorded on a Jasco J-1500 spectropolarimeter using a 0.5-mm pathlength quartz cuvette over the spectral range of 190–280 nm (scanning speed, 10 nm min^−1^; response time, 8 s; data pitch, 0.5 nm; spectral resolution, 1 nm; two accumulations). Protein samples were prepared at a concentration of 0.2 mg mL^−1^ in phosphate-buffered saline (PBS) and measured at room temperature. Thermal denaturation profiles of Ap3, Fd3, Rn1, and Random10E were recorded in PBS from 10 °C to 90 °C in 5 °C increments using the same spectropolarimeter settings and cuvette. For pH-dependent measurements, spectra were collected in a universal buffer composed of 20 mM sodium acetate, 20 mM Tris-HCl, and 20 mM BIS-Tris adjusted to pH 4–8. Guanidinium hydrochloride titrations were performed using a 6 M stock solution diluted to the desired final concentrations. Halophilic stability assays were conducted in PBS supplemented with NaCl to a final concentration of 1 M (from a 5 M NaCl stock), using identical acquisition parameters. All spectra were baseline-corrected, and the final data were expressed as mean residue molar ellipticity (θ) in deg·cm^2^·dmol^−1^·residue^−1^.

### Fold Robustness Simulations

Analyses of fold robustness were performed on selected sequences and their respective.pdb files. For wild-type structures, the file for the respective template structures was downloaded from the PDB database. Only the relevant part that was used for the re-design was kept matching the size of the designed version. For the ancient 10 alphabet designs, their structure, determined with NMR, was used. For the full 20 designed alphabet counterparts, ESMFold prediction (esmfold_v1() pretrained model) was used. Structural predictions of the mutated sequences were performed during simulations using the same ESMFold model as above. TM-Score was calculated with the TM-align package^49^. Mutations considered were substitutions only, randomly selected in every step of the simulation. The whole simulation process with both alphabets was repeated 20 times for each protein. The chance of introducing a specific amino acid was equal for all the options included within the alphabet. The number of introduced mutations was increasing by one until the full length of the protein was reached. For the selected 10 and full 20 amino acid alphabets, *mutations_simulation_10E*.*py* and *mutations_simulation_20F*.*py* scripts were used, respectively. All values and plots for TM-score vs the number of introduced mutations for all runs and all proteins can be found in the Supplementary data. To plot the data, the first occurrence of TM-score < 0.5 was found in every run, and the number of mutations was then plotted for each run for simulation. Statistical significance of the differences between simulations was calculated for the pairs of the same protein using the two alphabets and between all combinations of proteins (wild-type, designed with 10 and designed with 20) for simulations run with one alphabet. Mann-Whitney U test within the scipy package^50^ was used for all comparisons. The Seaborn package was used for graphical visualization ^51^. The differences were considered significant when p-value < 0.01. Tables with values for all comparisons can be found in the Supplementary data.

### NMR spectroscopy

NMR spectra were acquired at 25°C using an 850 MHz Bruker Avance HD III spectrometer, equipped with a triple-resonance (^15^N/^13^C/1H) cryoprobe. The sample volume was 0.35 ml, 50 mM NaH2PO4, pH 6.5, 280 mM NaCl, 5% D2O/95% H2O. A series of double- (^1^H-^15^N HSQC, ^1^H-^13^C HSQC) and triple-resonance spectra (HNCO, HN(CA)CO, HNCACB, CBCA(CO)NH, HBHA(CO)NH, (H)CC(CO)NH, HCCH-TOCSY from standard Bruker pulse program library were recorded to obtain sequence-specific resonance assignment. in POKY^52^. 1H–1H distance restraints were derived from the 3D ^15^N/^1^H NOESY-HSQC and ^13^C/^1^H NOESY-HMQC spectra using a 100 ms mixing time. Structure determination procedure followed a well-established protocol as described previously^53^. The structural calculation was carried out in CYANA^54^ using NOESY data in combination with backbone torsion angle restraints, generated from assigned chemical shifts using the program TALOS+^55^. The automated NOE assignment and structure determination protocol was used for automatic NOE cross-peak assignment. Subsequently, five cycles of simulated annealing combined with redundant dihedral angle restraints were used to calculate a set of converged structures with no significant restraint violations (distance and van der Waals violations < 0.5Å and dihedral angle constraint violations < 5°). The 30 structures with the least restraint violations were further refined in explicit solvent using YASARA^56^ and subjected to further analysis using the Protein Structure Validation Software suite (www.nesg.org). The statistics for the resulting structures are summarized in Table S1-3. The structures, NMR restraints and resonance assignments were deposited in the Protein Data Bank (PDB ID 9SGV, 9SGJ, 9SGW and BMRB (accession code: 53280, 53283, 35014) for Ap3, Fd3 and Rn2, respectively.

## Acknowledgments

We thank Dr. Radko Souček for performing the amino acid analyses and Filip Buchel for assistance with library FACS sorting.

## Funding

This work was supported by a Research Grant from HFSP https://doi.org/10.52044/HFSP.RGEC272023.pc.gr.168579.

Primus grant PRIMUS/20/SCI/012 from Charles University.

The authors acknowledge the Imaging Methods Core Facility at BIOCEV, an institution supported by the MEYS CR (LM2023050 Czech-BioImaging) for their support & assistance in this work, and particularly Ondrej Honc.

CMS CF Biophysical techniques of CIISB, Instruct-CZ Centre, supported by MEYS CR Infrastructure project LM2018127 and European Regional Development Fund-Project, “UP CIISB” (No. CZ.02.1.01/0.0/0.0/18_046/0015974).

Computational resources for the mutational tolerance analyses were provided by the e-INFRA CZ project (ID: 90254), supported by the Ministry of Education, Youth and Sports of the Czech Republic.

This work was partially supported by the Wallenberg AI, Autonomous Systems and Software Program (WASP) funded by the Knut and Alice Wallenberg Foundation.

## Author contributions

Conceptualization: KH, IA

Methodology: KH, IA, VGG, SA, VV, PS, TN

Investigation: VGG, SA, PS, ZR, TN, LB, SP, JM, AK

Visualization: VGG, SA, VV, PS, TN

Funding acquisition: KH, IA

Project administration: KH, IA

Supervision: KH, IA, VV, VGG

Writing – original draft: KH, IA, VGG, SA, VV, TN

Writing – review & editing: KH, IA, VGG, SA

## Competing interests

The authors declare that they have no competing interests.

## Data and materials availability

All original code developed for this study and raw data is available at https://github.com/SIAndersson/AncientProteinScripts. The structures, NMR restraints and resonance assignments were deposited in the Protein Data Bank (PDB ID 9SGV, 9SGJ, 9SGW and BMRB (accession code: 53280, 53283, 35014) for Ap3, Fd3 and Rn2, respectively. All other data needed to evaluate the conclusions are available in the manuscript and supplementary materials.

## Supplementary Materials

Supplementary Text S1

Figs. S1 to S11

Tables S1 to S3

## Notes

### Competing Interest Statement

The authors have declared no competing interest.

## References and Notes

1. Brooks, D. J., Fresco, J. R., Lesk, A. M. & Singh, M. Evolution of Amino Acid Frequencies in Proteins Over Deep Time: Inferred Order of Introduction of Amino Acids into the Genetic Code. Molecular Biology and Evolution 19, 1645–1655 (2002).

2. Crapitto, A. J., Campbell, A., Harris, A. & Goldman, A. D. A consensus view of the proteome of the last universal common ancestor. Ecology and Evolution 12, e8930 (2022).

3. Philip, G. K. & Freeland, S. J. Did Evolution Select a Nonrandom “Alphabet” of Amino Acids? Astrobiology 11, 235–240 (2011).

4. Weber, A. L. & Miller, S. L. Reasons for the occurrence of the twenty coded protein amino acids. J Mol Evol 17, 273–284 (1981).

5. Fried, S. D., Fujishima, K., Makarov, M., Cherepashuk, I. & Hlouchova, K. Peptides before and during the nucleotide world: an origins story emphasizing cooperation between proteins and nucleic acids. J. R. Soc. Interface. 19, 20210641 (2022).

6. Stephenson, J. D. & Freeland, S. J. Unearthing the Root of Amino Acid Similarity. J Mol Evol 77, 159–169 (2013).

7. Higgs, P. G. & Pudritz, R. E. A Thermodynamic Basis for Prebiotic Amino Acid Synthesis and the Nature of the First Genetic Code. Astrobiology 9, 483–490 (2009).

8. Murphy, L. R., Wallqvist, A. & Levy, R. M. Simplified amino acid alphabets for protein fold recognition and implications for folding. Protein Engineering, Design and Selection 13, 149– 152 (2000).

9. Solis, A. D. Reduced alphabet of prebiotic amino acids optimally encodes the conformational space of diverse extant protein folds. BMC Evol Biol 19, 158 (2019).

10. Tretyachenko, V. et al. Modern and prebiotic amino acids support distinct structural profiles in proteins. Open Biol. 12, 220040 (2022).

11. Makarov, M. et al. Early Selection of the Amino Acid Alphabet Was Adaptively Shaped by Biophysical Constraints of Foldability. J. Am. Chem. Soc. 145, 5320–5329 (2023).

12. Longo, L. M. et al. Primordial emergence of a nucleic acid-binding protein via phase separation and statistical ornithine-to-arginine conversion. Proc. Natl. Acad. Sci. U.S.A. 117, 15731–15739 (2020).

13. Giacobelli, V. G. et al. In Vitro Evolution Reveals Noncationic Protein–RNA Interaction Mediated by Metal Ions. Molecular Biology and Evolution 39, msac032 (2022).

14. Longo, L. M., Tenorio, C. A., Kumru, O. S., Middaugh, C. R. & Blaber, M. A single aromatic core mutation converts a designed “primitive” protein from halophile to mesophile folding. Protein Science 24, 27–37 (2015).

15. Longo, L. M., Lee, J. & Blaber, M. Simplified protein design biased for prebiotic amino acids yields a foldable, halophilic protein. Proc. Natl. Acad. Sci. U.S.A. 110, 2135–2139 (2013).

16. Longo, L. M. & Blaber, M. Protein design at the interface of the pre-biotic and biotic worlds. Archives of Biochemistry and Biophysics 526, 16–21 (2012).

17. Tanaka, J., Doi, N., Takashima, H. & Yanagawa, H. Comparative characterization of random‐sequence proteins consisting of 5, 12, and 20 kinds of amino acids. Protein Science 19, 786– 795 (2010).

18. Newton, M. S., Morrone, D. J., Lee, K. & Seelig, B. Genetic Code Evolution Investigated through the Synthesis and Characterisation of Proteins from Reduced‐Alphabet Libraries. ChemBioChem 20, 846–856 (2019).

19. Zhao, F. & Akanuma, S. Ancestral Sequence Reconstruction of the Ribosomal Protein uS8 and Reduction of Amino Acid Usage to a Smaller Alphabet. J Mol Evol 91, 10–23 (2023).

20. Walter, K. U., Vamvaca, K. & Hilvert, D. An Active Enzyme Constructed from a 9-Amino Acid Alphabet. Journal of Biological Chemistry 280, 37742–37746 (2005).

21. Akanuma, S., Kigawa, T. & Yokoyama, S. Combinatorial mutagenesis to restrict amino acid usage in an enzyme to a reduced set. Proc. Natl. Acad. Sci. U.S.A. 99, 13549–13553 (2002).

22. Shibue, R. et al. Comprehensive reduction of amino acid set in a protein suggests the importance of prebiotic amino acids for stable proteins. Sci Rep 8, 1227 (2018).

23. Makarov, M. et al. Enzyme catalysis prior to aromatic residues: Reverse engineering of a dephospho‐CoA kinase. Protein Science 30, 1022–1034 (2021).

24. Caetano-Anollés, G., Kim, H. S. & Mittenthal, J. E. The origin of modern metabolic networks inferred from phylogenomic analysis of protein architecture. Proc. Natl. Acad. Sci. U.S.A. 104, 9358–9363 (2007).

25. Chandonia, J.-M. et al. SCOPe: improvements to the structural classification of proteins – extended database to facilitate variant interpretation and machine learning. Nucleic Acids Research 50, D553–D559 (2022).

26. Watson, J. L. et al. De novo design of protein structure and function with RFdiffusion. Nature 620, 1089–1100 (2023).

27. Dauparas, J. et al. Robust deep learning–based protein sequence design using ProteinMPNN. Science 378, 49–56 (2022).

28. Lin, Z. et al. Evolutionary-scale prediction of atomic-level protein structure with a language model. Science 379, 1123–1130 (2023).

29. Jumper, J. et al. Highly accurate protein structure prediction with AlphaFold. Nature 596, 583– 589 (2021).

30. Van Kempen, M. et al. Fast and accurate protein structure search with Foldseek. Nat Biotechnol 42, 243–246 (2024).

31. Sillitoe, I. et al. CATH: increased structural coverage of functional space. Nucleic Acids Research 49, D266–D273 (2021).

32. Ieremie, I., Ewing, R. M. & Niranjan, M. Protein language models meet reduced amino acid alphabets. Bioinformatics 40, btae061 (2024).

33. Hlouchová, K. Peptides En Route from Prebiotic to Biotic Catalysis. Acc. Chem. Res. 57, 2027–2037 (2024).

34. Milner-White, E. J. & Russell, M. J. Functional Capabilities of the Earliest Peptides and the Emergence of Life. Genes 2, 671–688 (2011).

35. Van Der Gulik, P., Massar, S., Gilis, D., Buhrman, H. & Rooman, M. The first peptides: The evolutionary transition between prebiotic amino acids and early proteins. Journal of Theoretical Biology 261, 531–539 (2009).

36. Xu, J. & Zhang, Y. How significant is a protein structure similarity with TM-score = 0.5? Bioinformatics 26, 889–895 (2010).

37. Magis, C. et al. Structure-Based Secondary Structure-Independent Approach To Design Protein Ligands: Application to the Design of Kv1.2 Potassium Channel Blockers. J. Am. Chem. Soc. 128, 16190–16205 (2006).

38. Zaia, D. A. M., Zaia, C.T.B.V. & De Santana, H. Which Amino Acids Should Be Used in Prebiotic Chemistry Studies? Orig Life Evol Biosph 38, 469–488 (2008).

39. Ilardo, M. et al. Adaptive Properties of the Genetically Encoded Amino Acid Alphabet Are Inherited from Its Subsets. Sci Rep 9, 12468 (2019).

40. Koga, R. et al. Robust folding of a de novo designed ideal protein even with most of the core mutated to valine. Proc. Natl. Acad. Sci. U.S.A. 117, 31149–31156 (2020).

41. Bloom, J. D., Labthavikul, S. T., Otey, C. R. & Arnold, F. H. Protein stability promotes evolvability. Proc. Natl. Acad. Sci. U.S.A. 103, 5869–5874 (2006).

42. Bloom, J. D. et al. Evolution favors protein mutational robustness in sufficiently large populations. BMC Biol 5, 29 (2007).

43. Rothschild, L. J. et al. Building Synthetic Cells─From the Technology Infrastructure to Cellular Entities. ACS Synth. Biol. 13, 974–997 (2024).

44. Chen, S., Zhou, Y., Chen, Y. & Gu, J. fastp: an ultra-fast all-in-one FASTQ preprocessor. Bioinformatics 34, i884–i890 (2018).

45. Fu, L., Niu, B., Zhu, Z., Wu, S. & Li, W. CD-HIT: accelerated for clustering the next-generation sequencing data. Bioinformatics 28, 3150–3152 (2012).

46. Li, H. & Durbin, R. Fast and accurate short read alignment with Burrows–Wheeler transform. Bioinformatics 25, 1754–1760 (2009).

47. Li, H. et al. The Sequence Alignment/Map format and SAMtools. Bioinformatics 25, 2078– 2079 (2009).

48. Anders, S., Pyl, P. T. & Huber, W. HTSeq—a Python framework to work with high-throughput sequencing data. Bioinformatics 31, 166–169 (2015).

49. Zhang, Y. TM-align: a protein structure alignment algorithm based on the TM-score. Nucleic Acids Research 33, 2302–2309 (2005).

50. Virtanen, P. et al. SciPy 1.0: fundamental algorithms for scientific computing in Python. Nat Methods 17, 261–272 (2020).

51. Waskom, M. seaborn: statistical data visualization. JOSS 6, 3021 (2021).

52. Lee, W., Rahimi, M., Lee, Y. & Chiu, A. POKY: a software suite for multidimensional NMR and 3D structure calculation of biomolecules. Bioinformatics 37, 3041–3042 (2021).

53. Cermakova, K. et al. A ubiquitous disordered protein interaction module orchestrates transcription elongation. Science 374, 1113–1121 (2021).

54. Güntert, P. & Buchner, L. Combined automated NOE assignment and structure calculation with CYANA. J Biomol NMR 62, 453–471 (2015).

55. Shen, Y., Delaglio, F., Cornilescu, G. & Bax, A. TALOS+: a hybrid method for predicting protein backbone torsion angles from NMR chemical shifts. J Biomol NMR 44, 213–223 (2009).

56. Harjes, E. et al. GTP-Ras Disrupts the Intramolecular Complex of C1 and RA Domains of Nore1. Structure 14, 881–888 (2006).

