## Supplementary materials for "Ancient amino acid sets enable stable protein folds"

Valerio Guido Giacobelli & Sofia Andersson *et al.,*

**The PDF file includes:**

Supplementary Text S1
Figs. S1 to S11
Table S1 to S3

**Supplementary Text**

Selected Ancient DNA and protein sequences

> Ap1
*ATGTCTACTGCTGCGGAAATCGAAGAGTTGTTGCTGGAAGCTGGCGCTGCGGGCGAGGC
TGTTACTGTGGCCGAAATTGCCGCCACGCTTGGCATTCCCGTCAGCACTGCTTCCGAGAT
CACGTCGGAGGTGCTGGCCGCCACAGGATTGGCTGCTGAAGTAACAGGTACTACCATCA
CCATCGACCTGGGC*
> Ap1
MSTAAEIEELLLEAGAAGEAVTVAEIAATLGIPVSTASEITSEVLAATGLAAEVTGTTITID
LG
> Ap2
*ATGGGATCAGCGGAGGCCGCGCTGGCCGAGGCGGAAGAAGCGATCTTAGCCTTATTGGC
AGCGGGGCCACTGACTGTCGAGGAGTTAGCAGAAGCCGTCGGGCTTGCGCCCGCGACTA
CCTCGGAGCTGTTAGCTGAGTTGGTTGAAGAAGGCGCAGTGGTCGTAGTGGCTGAAGAG
GCGGGGGTTGTCACATTGGCTTTGGCGACG*
> Ap2
MGSAEAALAEAEEAILALLAAGPLTVEELAEAVGLAPATTSELLAELVEEGAVVVVAEE
AGVVTLALAT
> Ap3
*ATGGAAACCGCCGAAACTGCTGAGGCCATTTTAGCTCTGGCAGCAGAGGGACCCCTTTCG
TTGGCCGAGATTGCGGAGGCACTTGGCTTACCGCTGCCGACCGTCAGCGAATTGGTGGC
TGAACTTGAAGCTGAAGGTCTGTTAGTAACTGCACCAGATGGCTCCGTTTCACTTGCGGCA*
> Ap3
METAETAEAILALAAEGPLSLAEIAEALGLPLPTVSELVAELEAEGLLVTAPDGSVSLAA
> Fd1
*ATGACACTGGTGGCTACAGCGGCTGACGAAGCCGCTGAAGCGGAAGTCGTAGAGGCACT
TGAGGCGGCAGGGGCTGAGGCAGTAGCGGTGGAAGGAGGGACTATTACGGTAACTGCT
GCTGATGAGGAAGCCACCCTTGAACTGCTGGAGGCACTGGAAGCTGAAGGACTGTTATTA
AGCGTTGCCGCAGCC*
> Fd1
TLVATAADEAAEAEVVEALEAAGAEAVAVEGGTITVTAADEEATLELLEALEAEGLLLS
VAAA
> Fd2
*ATGTCTACCACTTCAACCCTTACGGTAACGGCAGACGCGGAGACAACGGAAACCATCACG
GAGCTGGCAGCAGAGGCCGGAGCGACCTCCACAACCGCCACCGAAGACGGTGCAACTG
TTTTAACGGTAACTGGGGACGCCGCAACGATTGAAGCTGTGGCGGAAGCCATTGCAGCTC
TTGGAATCACCCCGGATTCTACCACC*
> Fd2
MSTTSTLTVTADAETTETITELAAEAGATSTTATEDGATVLTVTGDAATIEAVAEAIAAL
GITPDSTT
> Fd3
*ATGGCCACGACAATTACTATCAGTGGGGACGCCGCGCTTTTGGCTGAGGCTCTTGAGGAA
GCCGAAGCCTTATTGGCCGCGGGTGTTATTGACGCAGTAGAAGTAGTCGATGGGGCTCTT*

*GTAATTACAGTTGCGCCAGCAGACGCAGAGGCCGTCGCGGAGGAACTTGCTGAAGCCGT*
*ACCAGGAATTACGGTAGAGATTTCTGCA*
>Fd3
MATTITISGDAALLAEALEEAEALLAAGVIDAVEVVDGALVITVAPADAEAVAEELAEA
VPGITVEISA
> Fd3 TEV
*ATGGCCACGACAATTACTATCAGTGGGGACGCCGCGCTTTTGGCTGAGGCTCTTGAGGAA*
*GCCGAAGCCTTATTGGCCGCGGGTGTTATTGACGCAGTAGAAGTAGTCGATGGGGCTCTT*
*GTAATTACAGTTGCGCCAGCAGACGCAGAGGCCGTCGCGGAGGAACTTGCTGAAGCCGT*
*ACCAGGAATTACGGTAGAGATTTCTGCAGAAAATCTGTACTTTCAGTCT*
> Fd3 TEV
MATTITISGDAALLAEALEEAEALLAAGVIDAVEVVDGALVITVAPADAEAVAEELAEA
VPGITVEISAENLYFQS
> Rn1
*ATGACTATCGTCACAGTGAGTAGCGCGGGAACAGGAACAACCCTGGACGGTACTGAAGTT*
*TCGCTGGAGGAAGCGGCGGCTGCGGACTTAATCGTGGCTGACACGGAGGCTGTGGCAG*
*CGGAACTTGCCGCGCTTGGGCTGCCTGTCGTTTTAGCTTCAGAGTTGGCAGAT*
> Rn1
MTIVTVSSAGTGTTLDGTEVSLEEAAAADLIVADTEAVAAELAALGLPVVLASELAD
> Rn2
*ATGACGGTTGTGACTATCACAGACACTACTCCGACAACAGCCACAGTAACAGTTTCGGAC*
*GCTGAAACTGGTGCTACCTTGGCCACTGCTGATGTTAGTACTGCTGATCCGGCTGAAGCT*
*GCCGAGGAAATCTTGGAGTTAGTCCTTGCTGTAGGCGCTACCGAGGTGACTGTAACGATC*
*ACCTTAGCCACTGCCGCTGAAGCCGCCGAGGCAGCTACAGAGTTGGCGGCTGCTCTTAC*
*CGAGCTGGCAGCAGAAGCTGGAATTACAGTCACTGTTACTGCTACTACAGCCGCTCCA*
> Rn2
MTVVTITDTTPTTATVTVSDAETGATLATADVSTADPAEAAEEILELVLAVGATEVTVTI
TLATAAEAAEAATELAAALTELAAEAGITVTVTATTAAP
> Rn3
*ATGACGTTAGCTCTTACGGTGGAGCTTGAAGGGGGAACAACGACGGTTACAGTCACTCTT*
*CCAGCCTCGGCCGGAGATACCACTCTTTCTACGACAGGTGAGGATTTAGCGGAGGCAGTA*
*ACTACGGCTTTGGAGGCGGCTGCTGAGGCCCTGGGGACTCCCGCAGATATTGCTGTGAC*
*TTTAAGTGTAGATGCAAGCGCCTCGGCCGAGGCCATCGCGGAGGCCGTGGCCACTATCA*
*CTGAAGCCGCTGCCGAGGCCGGTGCAACCAGCGTTACCGTAACAACTACAACTCTTGCT*
> Rn3
MTLALTVELEGGTTTVTVTLPASAGDTTLSTTGEDLAEAVTTALEAAAEALGTPADIAV
TLSVDASASAEAIAEAVATITEAAAEAGATSVTVTTTTLA

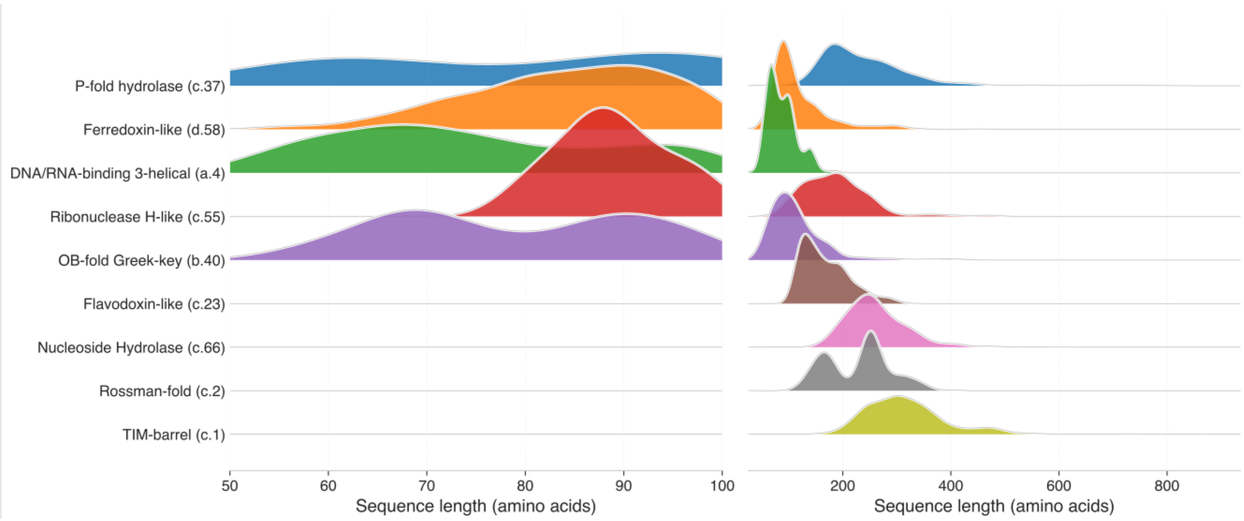


**Fig. S1. Comparison of protein length distributions across protein superfamilies.**
The right panel shows the distributions before truncation, and the left panel shows the distributions after restricting protein lengths to the range of 50–100 residues.

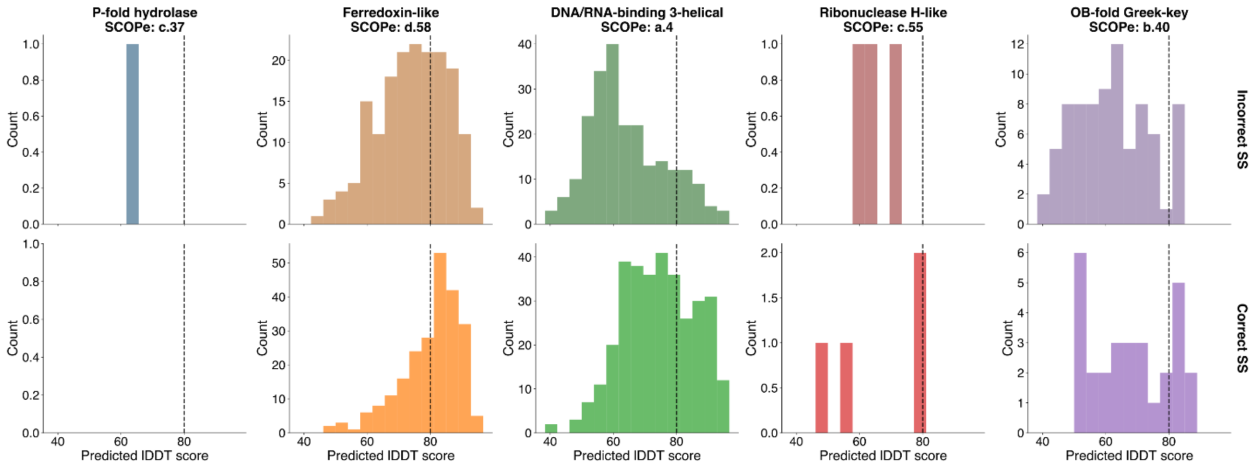


**Fig. S2. Comparison of pLDDT distributions across protein superfamilies.**

The top row displays designs with correct secondary structure, whereas the bottom row shows designs with incorrect secondary structure. The dashed line marks the minimum pLDDT threshold for acceptable designs (set to 80 in this analysis).

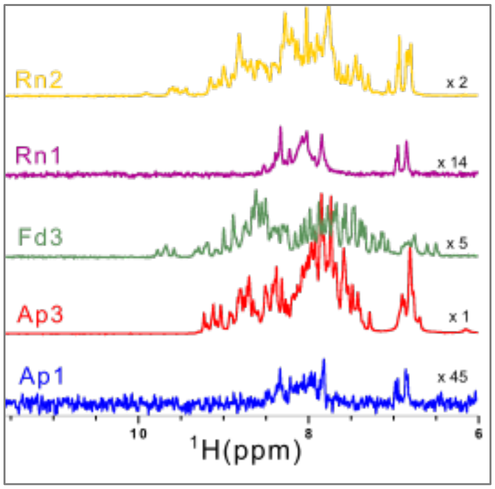


**Fig. S3. 1D NMR spectra of the selected ancient proteins.**
Comparison of 1D NMR spectra for Ap1, Ap3, Fd3, Rn1, and Rn2 in the 6–11.5 ppm region.

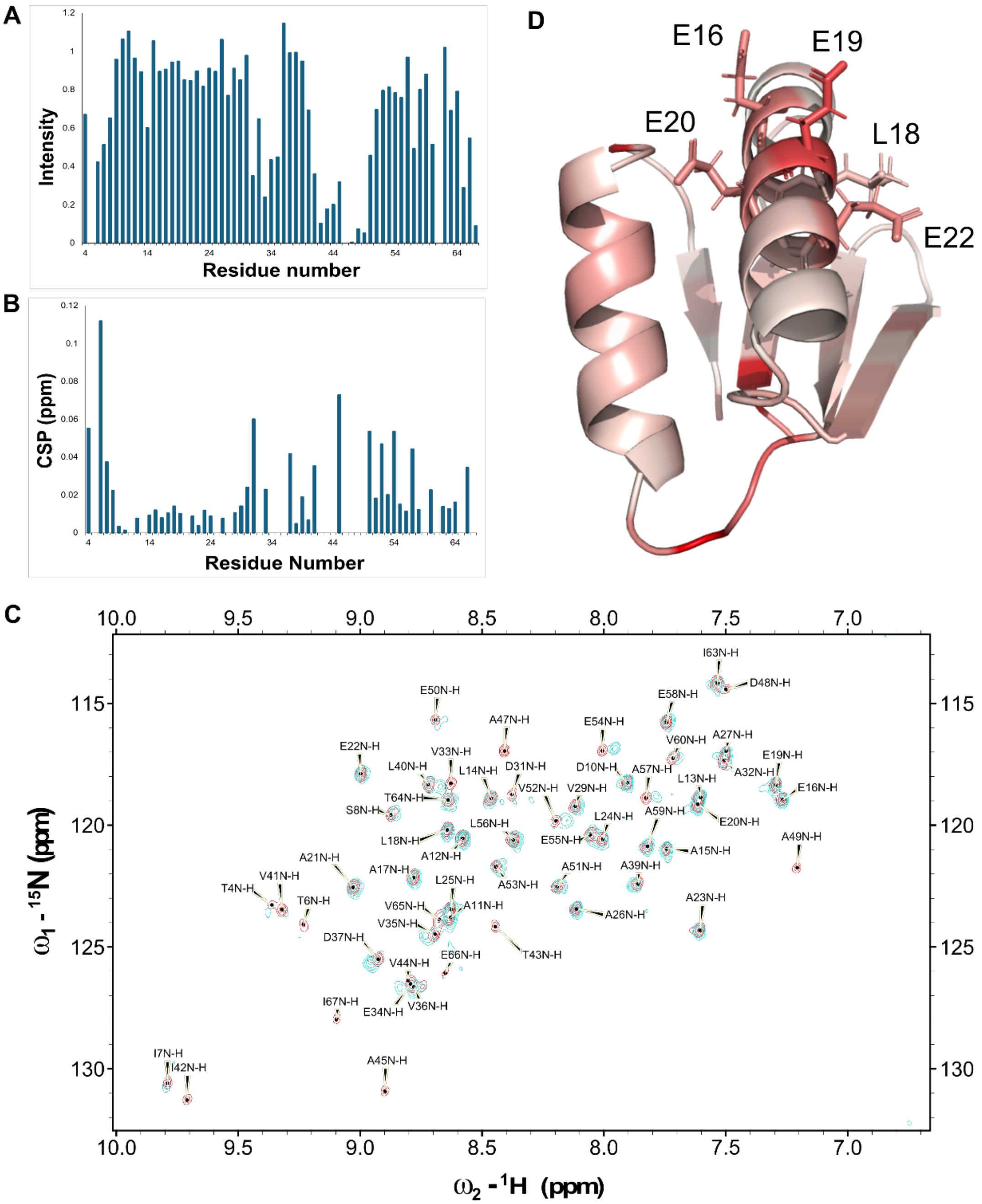


**Fig. S4. Copper-binding affinity assay of the Fd3 ancient protein.**

(A) Plot of the intensity ratio between the 2D NMR spectrum of Fd3 and that of Fd3 in the presence of $Cu^{2+}$. (B) Plot of the chemical shift perturbations between the 2D NMR spectrum of Fd3 and that of Fd3 with $Cu^{2+}$. (C) Overlay of the 2D NMR spectrum of Fd3 (blue peaks) and that of Fd3 with $Cu^{2+}$ at a molar ratio of 1:24 (red peaks). (D) Structure of Fd3 coloured according to the intensity ratio between the 2D NMR spectra of Fd3 and Fd3 with $Cu^{2+}$. Residues with smaller intensity ratios are shown in red. Residues involved in $Cu^{2+}$ binding are labelled.

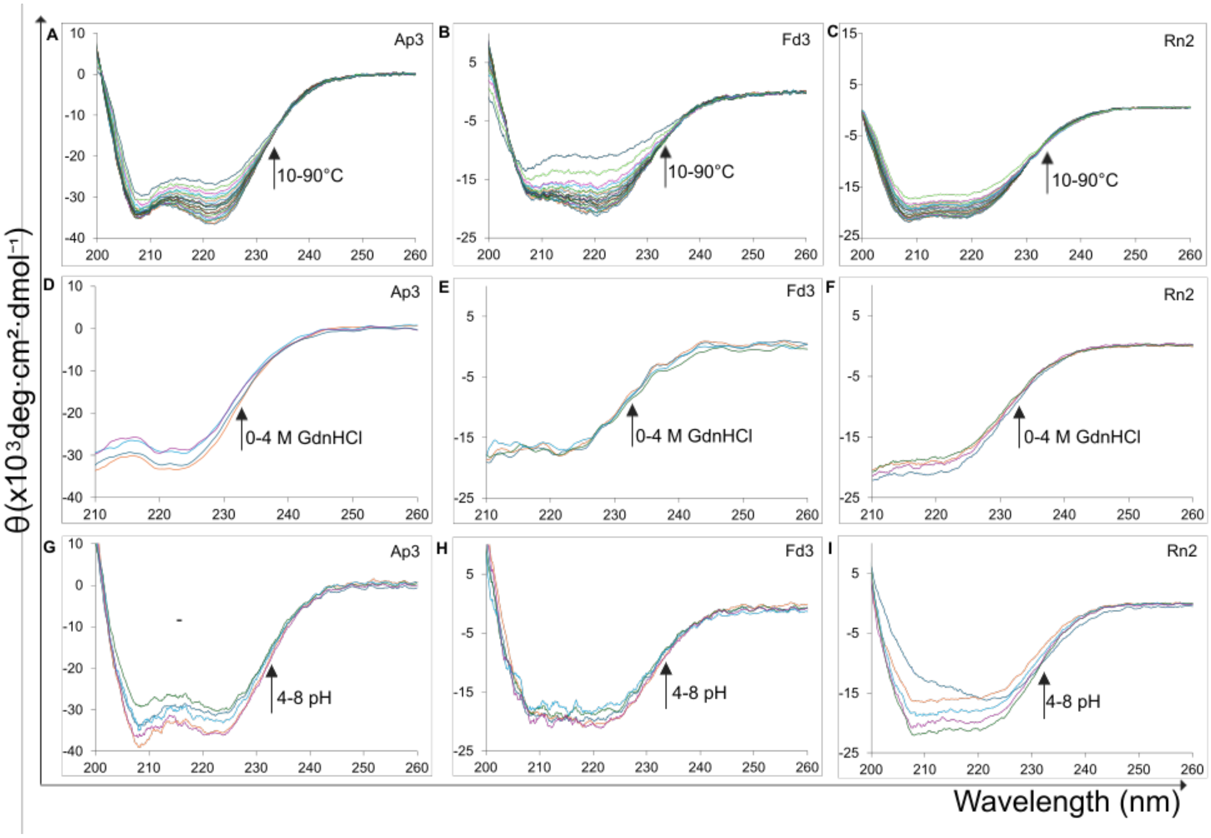


**Fig. S5. Temperature, chemical, and pH stability of selected ancient proteins.**

(A–C) CD spectra obtained over a temperature gradient from 10°C to 90°C in 5°C increments. (D–F) CD spectra during guanidine hydrochloride denaturation from 0 M to 4 M. (G–I) CD spectra over a pH range from 4 to 8.

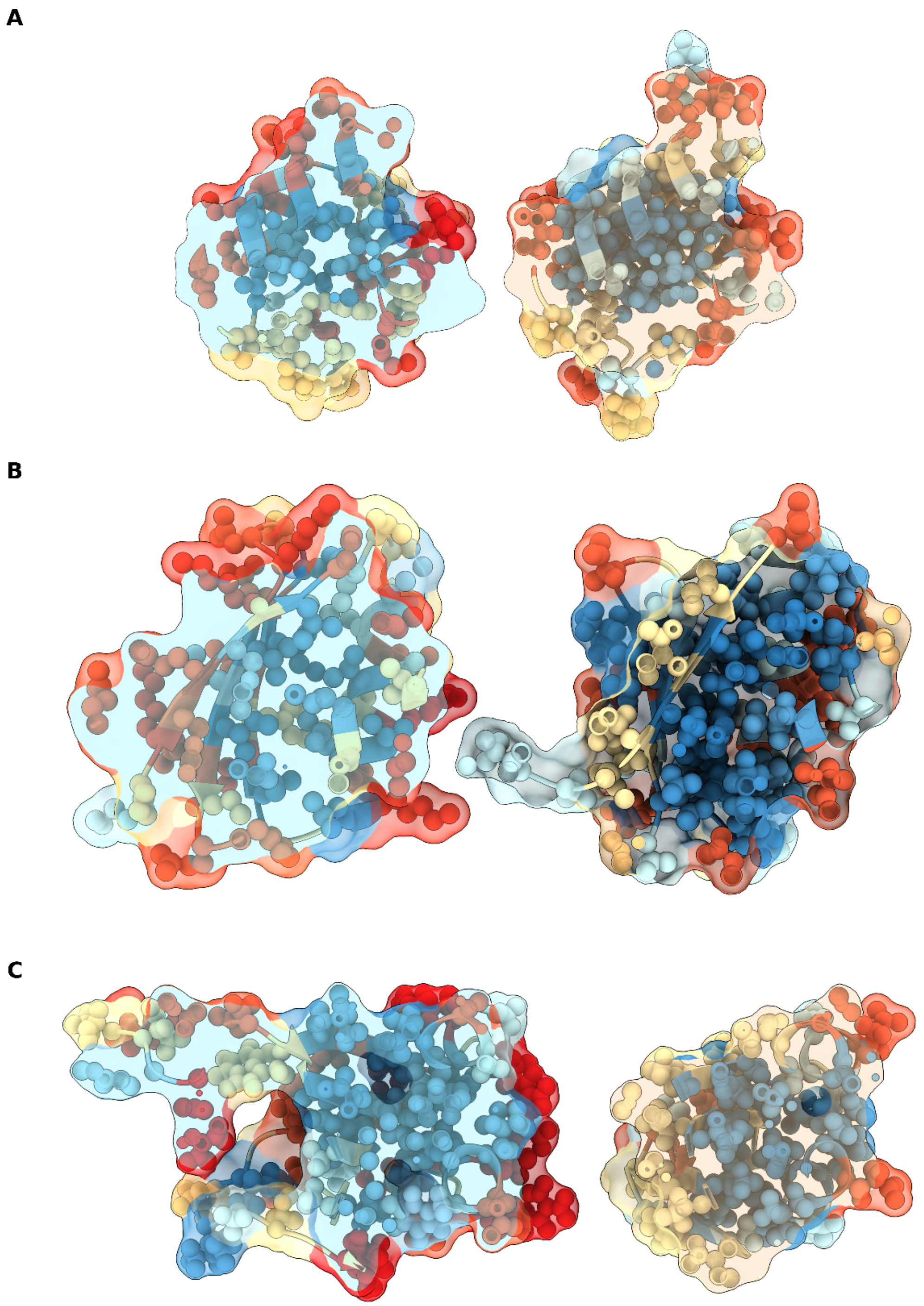


**Fig. S6. Comparison of hydrophobic cores between wild-type and designed protein NMR structures.**

In each panel, the wild-type structure is shown on the left and the designed structure on the right. Structures are colored according to the Kyte–Doolittle hydrophobicity scale, with blue indicating higher hydrophobicity and red indicating lower hydrophobicity. Approximately the central region of each protein is displayed. Panels show Ap3 (A), Fd3 (B), and Rn2 (C), respectively.

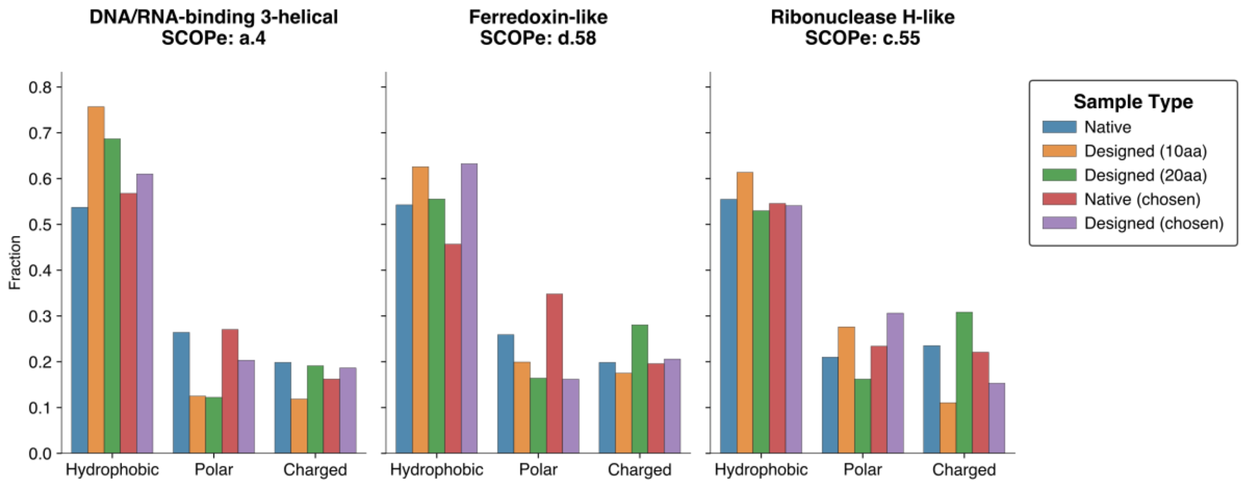


**Fig. S7. Comparison of biochemical properties of amino acids in native and designed proteins, for both full and restricted alphabets.**

The plots show fractions of different amino acid properties in the sequences of the native sequences, designed sequences, and the final three designs (Ap3, Fd3, and Rn2, here referred to as *chosen*). Only the sequences that ended in successful predictions were included in the comparison.

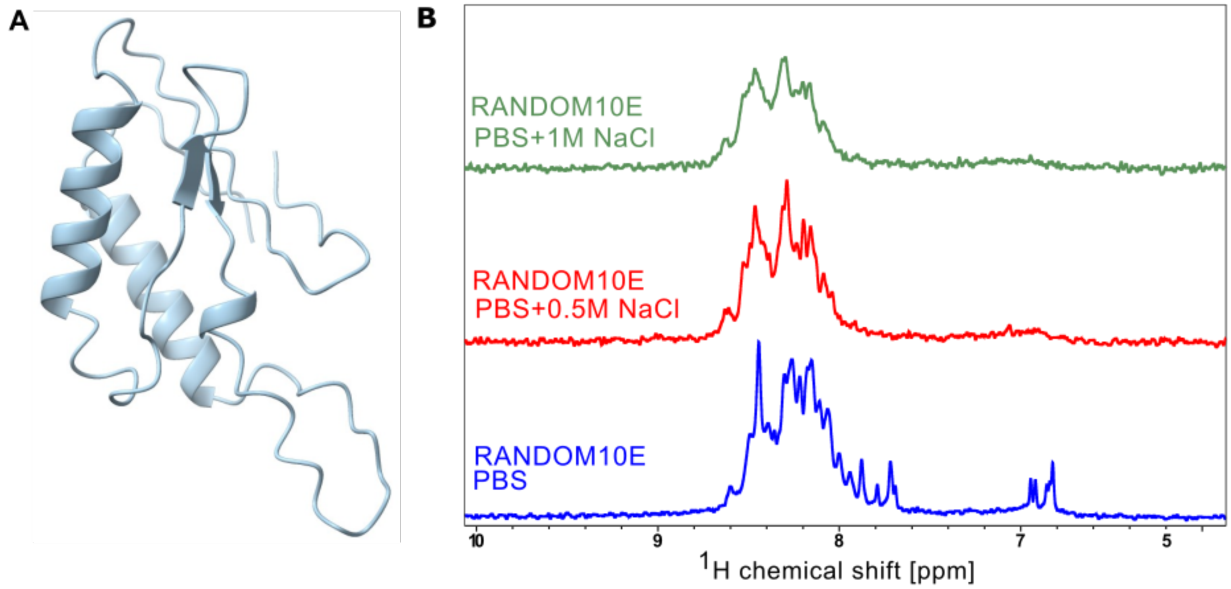


**Fig. S8. Structural and biophysical characterization of the Random 10E protein.**

(A) Random10E predicted structure generated using ESMFold. (B) 1D NMR spectra of Random 10E acquired in the presence of 0.5 M and 1 M NaCl.

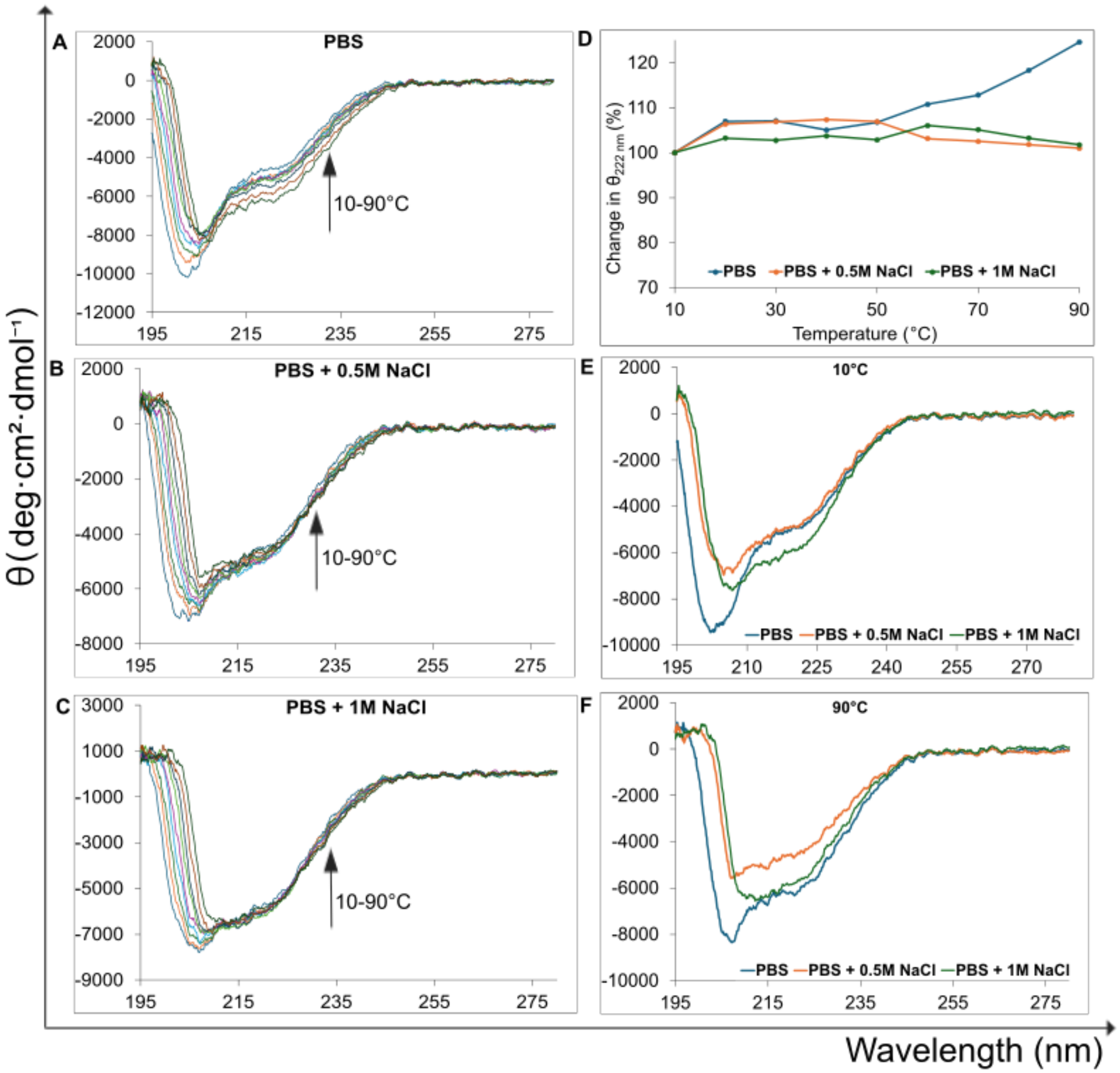


**Fig. S9. Temperature and NaCl stability of Random10E protein.**

(A–C) CD spectra obtained over a temperature gradient from 10°C to 90°C in 10 °C increments. D) Thermal stability of Random10E protein determined by monitoring the change of mean residue ellipticity, [θ] ($deg \cdot cm^2 \cdot dmol^{-1}$), at 222 nm as a function of temperature. (E-F) Comparison of CD spectra at different NaCl concentrations measured at 10°C and 90°C

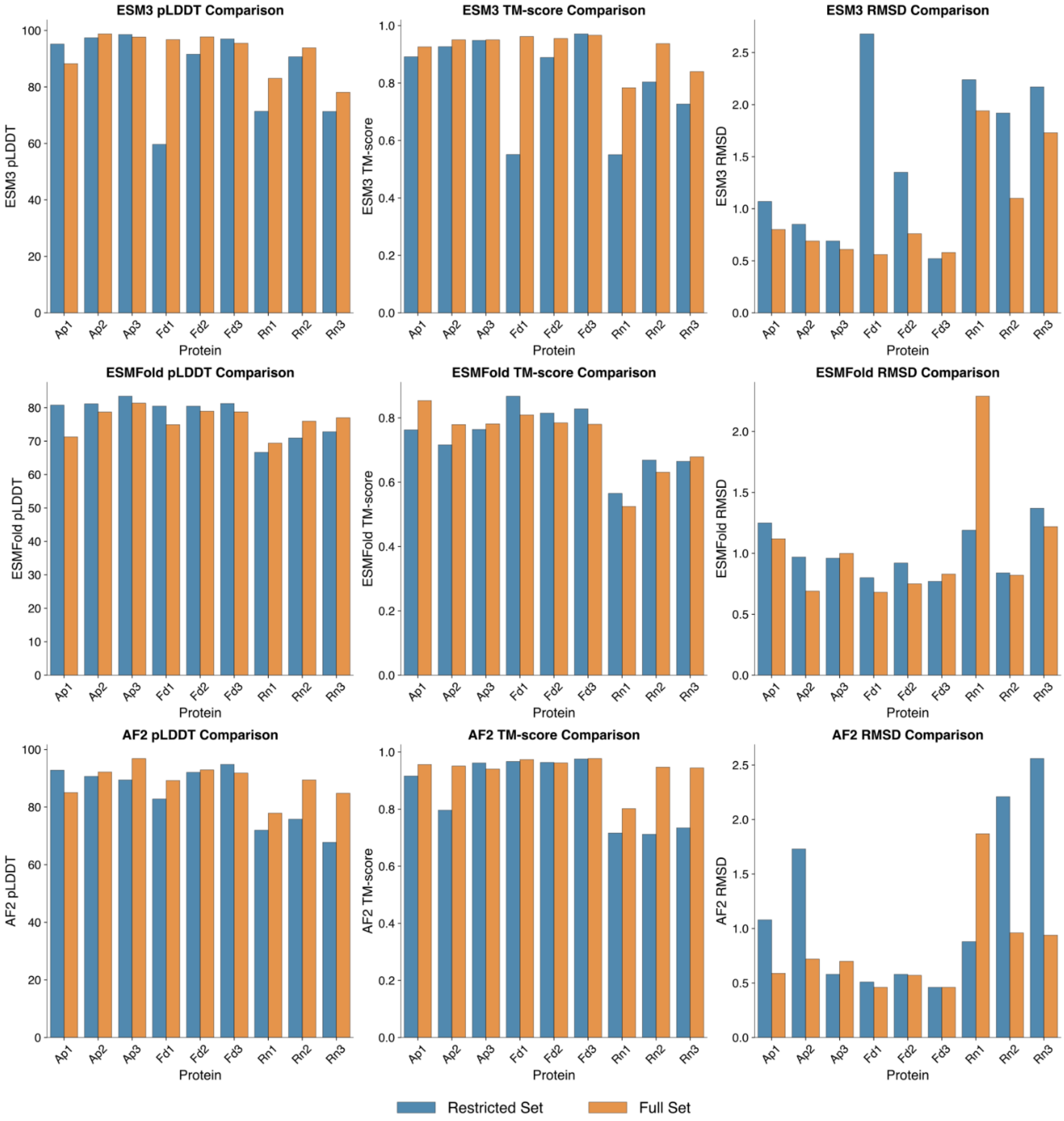


**Fig. S10. Comparison of metrics for full and restricted amino acid redesigns.**

Comparison of pLDDT, TM-score to design, and RMSD to design for the 9 final candidates using ESM3, ESMFold, and AlphaFold2.

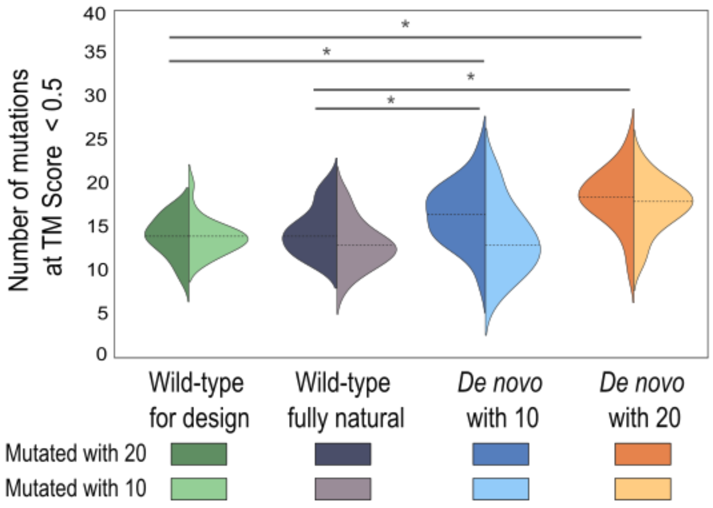


**Fig. S11. Mutational robustness of folds for Ap3 proteins with the addition of the fully natural wild-type sequence.**

Violin plots show the number of mutations required for the TM-score to fall below 0.5 (indicating loss of fold), based on 20 independent simulation runs per condition. The color legend below the plot specifies the amino acid alphabet used. Each half-violin represents a unique combination of amino acid alphabet and input protein. The dotted line denotes the median value. A significant difference was observed between the designed and fully natural wild-type proteins ($P \leq 0.01$, Mann–Whitney U test, indicated by *), whereas no significant difference was detected between the two wild-type structures.

| | |
|---|---|
| **Non-redundant distance and angle constrains** | |
| Total number of NOE restraints | 1291 |
| Intra-residue (i = j) | 221 |
| Sequential (\| i – j \| = 1) | 393 |
| Medium-range NOEs (1 < \| i – j \| < 5) | 356 |
| Long-range NOEs (\| i – j \| ≥ 5) | 321 |
| Torsion angles | 92 |
| Hydrogen bond restrains | 0 |
| Total number of restricting restraints | 1383 |
| Total restricting restraints per restrained residue | 24.3 |
| **Residual constraint violations** | |
| Distance violations per structure | |
| 0.1–0.2 Å | 1.9 |
| 0.2–0.5 Å | 0.37 |
| > 0.5 Å | 0 |
| r.m.s. of distance violation per constraint | 0.03 Å |
| Maximum distance violation | 0.0 Å |
| Dihedral angle violation per structure | |
| 1–10 ° | 1.03 |
| > 10 ° | 0 |
| r.m.s. of dihedral violations per constraint | 2.47 ° |
| **Ramachandran plot summary** | |
| Most favoured regions | 97.2% |
| Additionally allowed regions | 2.2% |
| Generously allowed regions | 0.6% |
| Disallowed regions | 0% |
| r.m.s.d. to the mean structure | all / ordered |
| All backbone atoms | 0.4 / 0.2 Å |
| All heavy atoms | 0.7 / 0.5 Å |
| **PDB entry** | 9SGV |
| **BMRB accession code** | 53280 |

**Table S1. Ap3 NMR Constraints and Statistics for the final set of structures.**

| | |
|---|---|
| **Non-redundant distance and angle constrains** | |
| Total number of NOE restraints | 777 |
| Intra-residue (i = j) | 219 |
| Sequential (\| i – j \| = 1) | 217 |
| Medium-range NOEs (1 < \| i – j \| < 5) | 141 |
| Long-range NOEs (\| i – j \| ≥ 5) | 200 |
| Torsion angles | 96 |
| Hydrogen bond restrains | 64 |
| Total number of restricting restraints | 937 |
| Total restricting restraints per restrained residue | 14.2 |
| **Residual constraint violations** | |
| Distance violations per structure | |
| 0.1–0.2 Å | 0.6 |
| 0.2–0.5 Å | 0.03 |
| > 0.5 Å | 0 |
| r.m.s. of distance violation per constraint | 0.04 Å |
| Maximum distance violation | 0.0 Å |
| Dihedral angle violation per structure | |
| 1–10 ° | 1.67 |
| > 10 ° | 0 |
| r.m.s. of dihedral violations per constraint | 2.77 ° |
| **Ramachandran plot summary** | |
| Most favoured regions | 96.5% |
| Additionally allowed regions | 3.5% |
| Generously allowed regions | 0.0% |
| Disallowed regions | 0.0% |
| r.m.s.d. to the mean structure | all / ordered |
| All backbone atoms | 1.9 / 0.3 Å |
| All heavy atoms | 2.5 / 0.6 Å |
| **PDB entry** | 9SGJ |
| **BMRB accession code** | 53283 |

**Table S2. Fd3 NMR Constraints and Statistics for the final set of structures.**

| | |
|---|---|
| **Non-redundant distance and angle constrains** | |
| Total number of NOE restraints | 1432 |
| Intra-residue (i = j) | 286 |
| Sequential (\| i – j \| = 1) | 504 |
| Medium-range NOEs (1 < \| i – j \| < 5) | 263 |
| Long-range NOEs (\| i – j \| ≥ 5) | 379 |
| Torsion angles | 162 |
| Hydrogen bond restrains | 84 |
| Total number of restricting restraints | 1678 |
| Total restricting restraints per restrained residue | 17.3 |
| **Residual constraint violations** | |
| Distance violations per structure | |
| 0.1–0.2 Å | 6.83 |
| 0.2–0.5 Å | 1.47 |
| > 0.5 Å | 0 |
| r.m.s. of distance violation per constraint | 0.06 Å |
| Maximum distance violation | 0.0 Å |
| Dihedral angle violation per structure | |
| 1–10 ° | 1.57 |
| > 10 ° | 0 |
| r.m.s. of dihedral violations per constraint | 2.71 ° |
| **Ramachandran plot summary** | |
| Most favoured regions | 95.8% |
| Additionally allowed regions | 4.2% |
| Generously allowed regions | 0.0% |
| Disallowed regions | 0% |
| r.m.s.d. to the mean structure | all / ordered |
| All backbone atoms | 4.1 / 0.3 Å |
| All heavy atoms | 4.4 / 0.5 Å |
| **PDB entry** | 9SGW |
| **BMRB accession code** | 35014 |

**Table S3. Rn2 NMR Constraints and Statistics for the final set of structures.**
